# Evolution takes multiple paths to evolvability when facing environmental change

**DOI:** 10.1101/2023.01.04.520634

**Authors:** Bhaskar Kumawat, Alexander Lalejini, Monica Acosta, Luis Zaman

**Affiliations:** Department of Ecology and Evolutionary Biology, University of Michigan, Ann Arbor, MI, 48109, USA; Center for the Study of Complex Systems, University of Michigan, Ann Arbor, MI, 48109, USA

**Keywords:** Digital Evolution, Evolvability, Mutation Rate

## Abstract

Living systems are surprisingly effective at exploiting new opportunities, as evidenced by the rapid emergence of antimicrobial resistance and novel pathogens. How populations attain this level of *evolvability* and the various ways it aids their survival are major open questions with direct implications for human health. Here, we use digital evolution to show that particular kinds of environments facilitate the simultaneous evolution of high mutation rates and a distribution of mutational effects skewed towards beneficial phenotypes. The evolved mutational neighborhoods allow rapid adaptation to previously encountered environments, whereas higher mutation rates aid adaptation to completely new environmental conditions. By precisely tracking evolving lineages and the phenotypes of their mutants, we show that evolving populations localize on phenotypic boundaries between distinct regions of genotype space. Our results demonstrate how evolution shapes multiple determinants of evolvability concurrently, fine-tuning a population’s adaptive responses to unpredictable or recurrent environmental shifts.

## Introduction

Biological life has flourished in every seemingly uninhabitable part of our world, and even as ongoing environmental change challenges ecosystems, rapid adaptation is a trademark response of natural populations. Emerging viral pathogens like SARS-CoV-2 and avian influenza (H5N1) are prime examples of populations that show a remarkable capacity to adapt to new hosts [1–5]. Similarly, controlling microbial growth is a constant struggle across industries—for example, in wastewater treatment, healthcare, and agriculture—precisely because antimicrobial resistance evolves so readily [6–8]. It is not obvious that living systems need to be so *evolvable* (i.e., able to generate adaptive variation), so why has evolution produced populations that are able to evolve so relentlessly? One challenge to addressing this question is that directly observing how evolvability evolves in natural populations is intractable due to the timescales and resolution of historical data that would be required [9–13]. A holistic picture of the processes that drive the evolution of evolvability is necessary to understand why evolution is so effective and to limit (or harness) its future potential [14].

To thrive in a rapidly changing world, organisms must give rise to mutant offspring that can survive impending environmental shifts [15–18]. At its core, Darwinian evolution is fueled by the continual emergence and fixation of new phenotypic variants in a population [19]. Such phenotypic variants arise as a result of two coupled processes. Replication of genetic material during the reproduction of an organism is susceptible to occasional errors that lead to mutations with a frequency quantified by the *mutation rate* [20]. However, only a small proportion of errors result in adaptive phenotypic change, since most mutations are neutral or deleterious [21]. Because of this property, mutation rates often rapidly decay, where they are maintained above zero through genetic drift [22, 23]. The set of mutants accessible from existing genotypes in a population and their corresponding phenotypes (i.e., the *mutational neighborhood*) governs the fraction of mutations that are ultimately beneficial [24]. Collectively, the mutation rate and mutational neighborhood govern the pace at which populations generate new phenotypes, providing the raw materials on which natural selection can act [12]. Nonetheless, the ability of populations to maintain high fitness becomes challenging in changing environments, where phenotypes can only be transiently well-adapted.

Prior work on phenotypic bias in mutations probes the ability of populations to adapt to specific environments [15, 16, 25–27]. In contrast, macroevolutionary studies view evolvability as a latent property of clades that promotes phenotypic divergence and diversification regardless of any selective advantage this variation provides [28, 29]. Alternatively, work on mutators and mutational modifiers focuses on an escalation in the rate at which variation is created, but ignores the phenotypic identity of the variants [30–32]. Bridging these scales, recent work suggests that microevolutionary evolvability shapes the rate of macroevolutionary diversification because the gradient of selection is constantly being perturbed as environments shift [33]. It is clear that the term evolvability is used ambiguously to describe several distinct processes in evolution. Furthermore, the conditions that shape the simultaneous evolution of multiple determinants of evolvability, how these determinants influence each other, and the way they aid adaptation in specific or novel environments remains unexplored [13].

In this work, we use populations of self-replicating computer programs in the digital evolution frame-work *Avida* to study how environmental conditions influence the evolution of evolvability [34]. The computational nature of this systems affords us unparalleled precision in tracking the mutational changes that occur along lineages over evolutionary time, while recording information about the external environment when mutations arise. In addition, it allows an in-depth analysis of the mutational landscapes of these lineages, extending to the second mutational step and encompassing over three million mutants per genotype. Using this framework, we study the mutation rates and structures of the mutational neighborhood that evolve as a result of exposure to different environments. We assess the performance of evolved organisms in their native and non-native conditions to highlight the way these determinants shape adaptation. Our findings demonstrate the existence of specific modes of environment change where the mutation rate and mutational neighborhoods evolve in ways that enhance future evolutionary innovation (Fig. 1). Taking advantage of tractability these *digital organisms* allows us to overcome the challenges previously limiting the direct observation of evolving evolvability.

**Figure 1:**
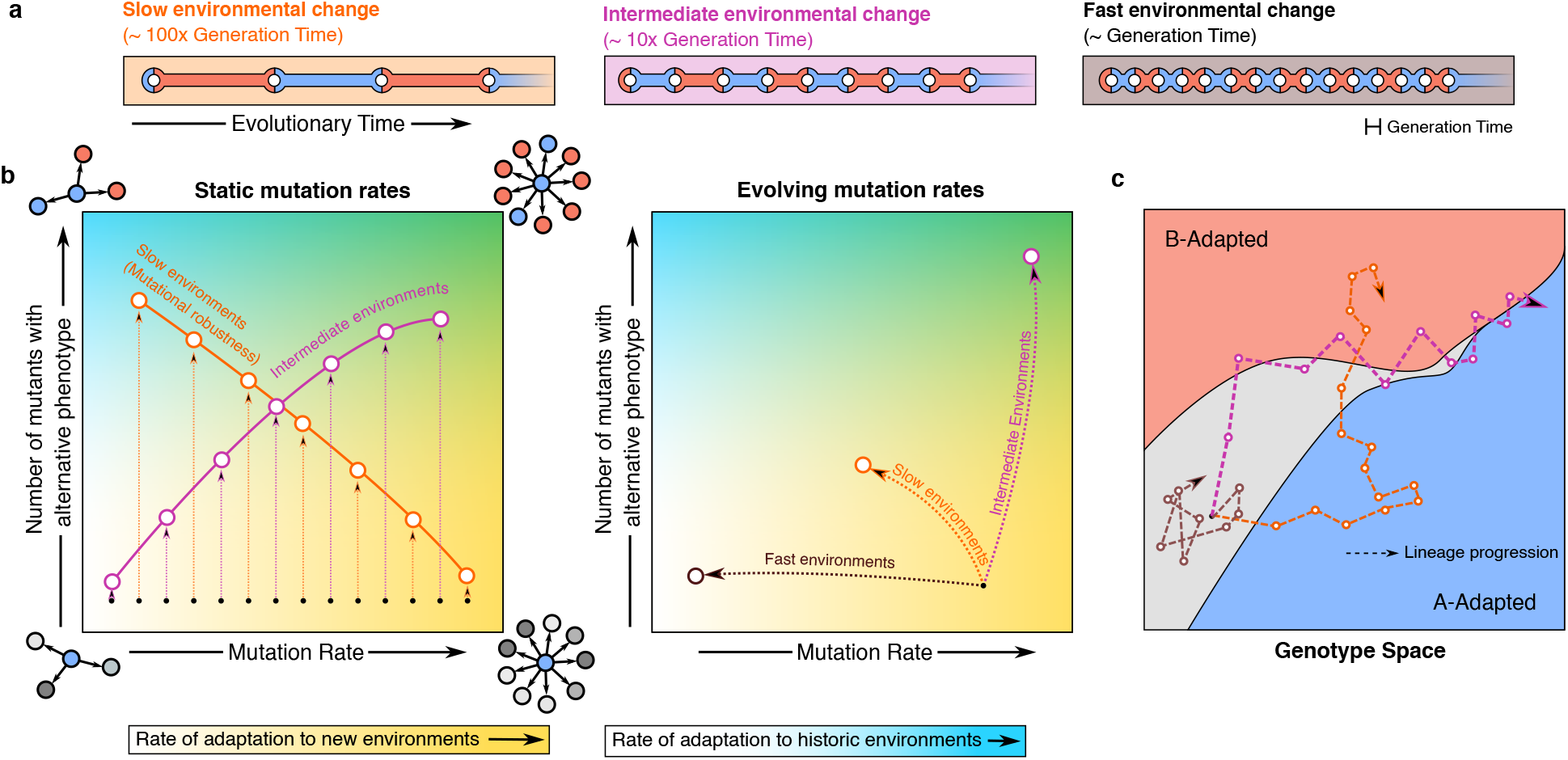
Schematic summarizing some key results for three environments. (**a**) We study evolution under environments that switch between two states (A and B) at three different rates. Evolution in all environments concludes in a specific environmental state (say A) and thus “alternate phenotype” here is the phenotype adapted to environment B. The switches in the three environments happen with an interval that is on the order of one generation, 10 generations, or 100 generations. (**b**) We perform two types of experiments, those where the mutation rate is fixed (left) and those where the mutation rates are allowed to evolve (right). When mutation rates are fixed and the environment switches slowly (orange curve, left), we observe classical mutation robustness, with increasing mutation rates leading to a decrease in the number of mutants with an alternate phenotype. However, at intermediate rates (purple curve, left) increasing mutation rate favors development of alternate phenotype mutants in the mutational neighborhood. When mutation rates are allowed to evolve, the intermediate environments lead to high mutation rates as well as a large number of alternative phenotypes in the mutational neighborhood. We find that while elevated mutation rates help adapt to completely new environments (blue shade), an increasing number of alternate phenotype mutants allows for quicker adaptation to historic environments (yellow shades). Thus, intermediate environments promote evolution of evolvability through two different pathways and buffer the population against different types of environmental change. (**c**) Upon analyzing lineage dynamics, we find that intermediate environments lead to populations localizing on phenotypic boundaries in the genotype space, where alternate mutants are easily accessible.

## Results

### Evolution of the mutational neighborhood

The *genotype-phenotype map* is a conceptual tool that describes the connection between an organism’s genetic code and its phenotypic traits [35]. Because there is redundancy in most genotype-phenotype maps, the local mutational neighborhood of organisms with the same phenotype may differ, making some of them more prone to expressing alternate phenotypes when mutated [24, 35–37]. First, we tested if the propensity of genotypes to generate mutants with alternate phenotypes can evolve and identified environmental conditions that favored this evolution.

Starting with a naive ancestral genotype, we evolved Avida populations under six different environment change regimes for approximately 30,000 generations (or equivalently 300,000 Avida updates, with an average generation time of 10 updates). We fixed the per-site copy mutation rate of the organisms in these experiments at 0.001, corresponding to an average of one mutation per 1,000 copied Avida instructions. Our experimental Avida world supported a maximum of 22,500 digital organisms. We explored stable environments in two regimes—*Const-A*, which is always in environment state *A*, and *Const-B*, which is always in environment state *B*. Additionally, four regimes—*Cyclic*, *Cyclic (Slow)*, *Cyclic (Fast)*, and *Random*—fluctuate between environmental states. The *Cyclic-*regimes switch between recurrent environmental states (*A → B → A → B…*) at different rates. The *Cyclic (Slow)*, *Cyclic*, and *Cyclic (Fast)* regimes stochastically switch states every 3, 30, and 300 generations on average, respectively. The *Random* regime switches randomly between one of 64 possible environmental states every 30 generations on average. Figure 2a shows a schematic detailing these environmental dynamics. Note that the environmental states *A* and *B* have strong fitness trade-offs between traits, with adaptations to environment *A* being deleterious in environment *B*, and vice versa.

**Figure 2:**
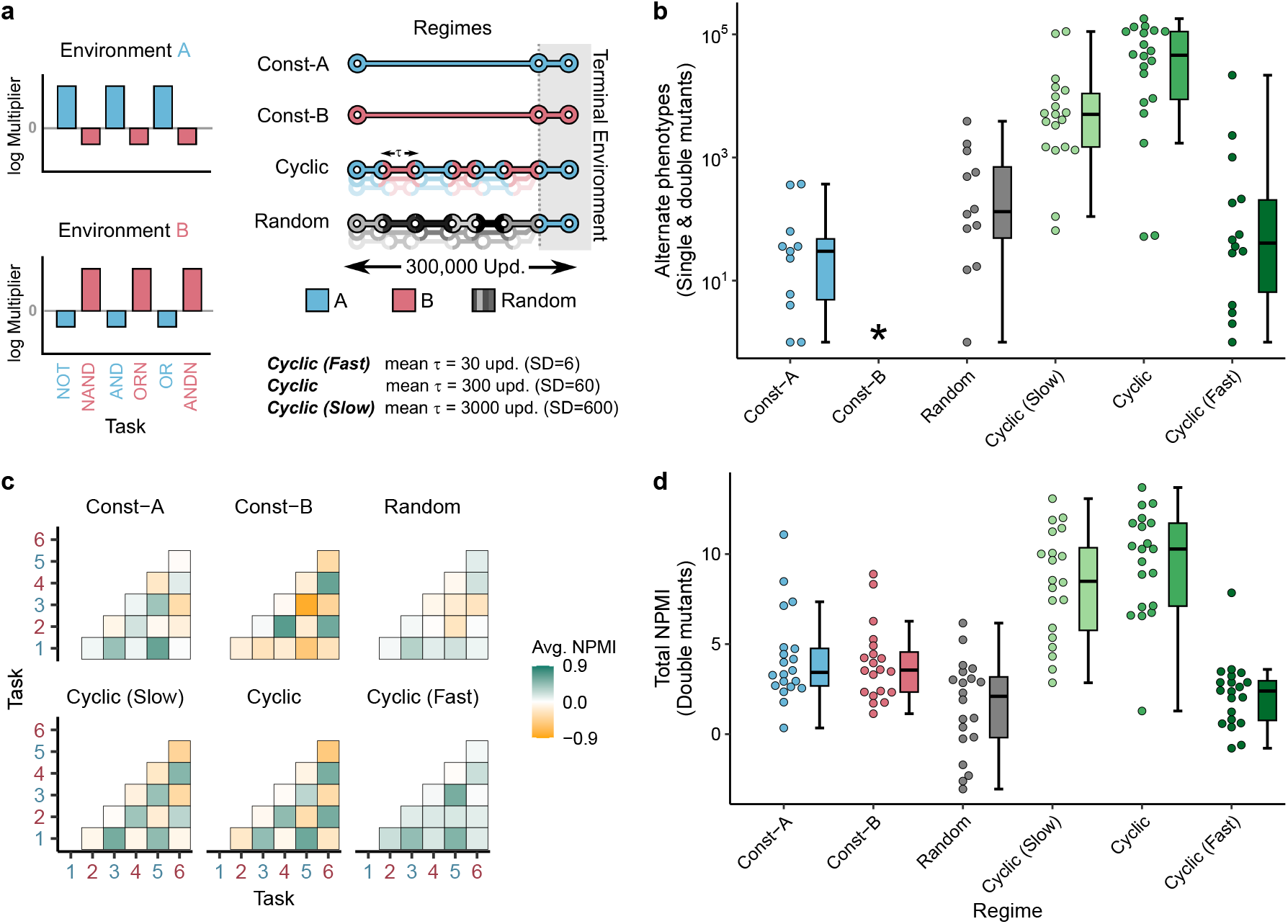
Evolution in a changing environment shapes the mutational neighborhood. (**a**) The two environments A and B recognize a set of six possible computational tasks performed by the organisms (left, x-axis). The fitness effects of task execution is determined by the respective fitness multipliers in each environment (see Methods). The two environments reward a complementary set of tasks. The schematic on the right details the different environmental change regimes used in this study and the environmental states they exhibit during evolution. The Cyclic-regimes switch between two environments with a waiting time (*τ*) picked from a Gaussian distribution, as detailed below the schematic. The *Random* regime rewards a randomly picked set of tasks and switches with a similar waiting time distribution as *Cyclic*. Each regime ends with 300 updates of evolution in a fixed terminal environment (grey region). (**b**) Total number of single and double mutants of the end-point dominant genotype with an alternate phenotype. Labels on the x-axis correspond to the six environmental regimes. (**c**) Normalized pointwise mutual information (NPMI) between task pairs for dominant genotypes isolated from populations evolved in different regimes (averaged over 20 replicates). The marginal and joint probabilities for calculating the NPMIs are measured using only the double mutants of the dominant genotype at the end of the experiment. Axes labels in blue and red denote tasks rewarded in environments A and B, respectively. (**d**) Total NPMI for the dominant genotypes isolated from the six environmental change regimes. Total NPMIs are calculated by summing up the individual task-pair NPMIs after assigning them a positive or negative sign based on their cognate or non-cognate nature. The boundaries of the boxplot represent the first (*Q*_1_) and third quartiles (*Q*_3_), with the median represented by the central horizontal line. Whiskers extend to maximum values within the upper (*Q*_3_ + 1.5 *× IQR*) and lower fences (*Q*_1_ − 1.5 *× IQR*), where *IQR* = *Q*_3_ *− Q*_1_.

Following experimental evolution, we isolated the dominant sequence (i.e., the genotype present at the highest frequency at the end of evolution) from each evolved population and created all possible single and double point mutations in those sequences (Fig. S1b). We then assayed the resulting phenotypes, counting the number of mutants with a phenotype that is exactly *alternative* to the phenotype of the evolved dominant genotype—if the dominant genotype was perfectly adapted to environment A, we counted the number of mutants that were adapted to environment B, and vice versa. Figure 2b shows the number of these mutants with the alternative phenotype that evolved under different regimes (Figure S2a details the overall distribution of all mutant adaptations).

Genotypes isolated from the *Cyclic* regime had a markedly larger pool of mutants expressing the alter-native phenotype when compared to genotypes from constant environments (median for *Cyclic*=45802.5 versus median for *Const-A*=1.0; Mann-Whitney *U* = 6, *n* = 20, *p* = 1.4 *×* 10*^−^*^7^, two-tailed. No alternate phenotype mutants were observed for *Const-B* genotypes). This escalation in alternate phenotypes was contingent on a predictable environment; the *Random* regime where future environmental states do not recapitulate historical conditions lacked genotypes with increased accessibility of alternate phenotypes (median for *Random*=16.0 versus median for *Const-A*= 1.0; Mann-Whitney *U* = 224, *n* = 20, *p* = 0.221, two-tailed). Notably, genotypes isolated from the intermediate *Cyclic* regime—with the environment switching every 30 generations on average—generated more alternate phenotypes on mutation compared to other regimes (median for *Cyclic (Slow)*=4556.0, median for *Cyclic*=45802.5; Mann-Whitney *U* = 312, *n* = 20, *p* = 0.0019, two-tailed). To test if these alternative phenotypes were enriched in the mutational neighborhood at the expense of robustness, we measured the correlation between the number of four different classes of mutants—alternate (perfectly adapted to the alternate environment), terminal (perfectly adapted to the terminal environment), intermediate (perfectly adapted to neither environments), and inviable (unable to self-replicate). The number of alternate phenotype mutants was not significantly correlated with changes in any other category (Fig. S2b). Thus, our observations support theoretical predictions that increased evolvability does not necessarily lead to a reduction in genetic robustness [27, 38].

These results indicate that the populations subjected to cyclically changing environments tend to evolve towards regions in the genotype space where adaptive phenotypes—those with high fitness in frequently encountered environments—are mutationally accessible. To explicitly test whether the mutational neighborhoods of evolved genotypes encode measurable information about the historic environment states encountered by the population, we analyzed the normalized pointwise mutual information (NPMI) between tasks. The NPMI metric calculates whether the co-occurrence of any two tasks within a genotype’s mutational neighborhood is significantly higher than expected assuming independent task probabilities (see Methods section). Figure 2c shows the NPMI values for different task pairs derived from genotypes evolved in all six regimes. We observed that the mutational neighborhood of organisms from changing environments systematically captured the pattern of tasks rewarded in different environmental states. Specifically, in *Cyclic* and *Cyclic (Slow)* regimes, task pairs that are rewarded concurrently in an environment state (i.e., are *cognate*) yielded positive NPMIs whereas task pairs that were rewarded in alternate environments had a negative NPMI.

We calculated the total NPMI by consolidating the task-pair NPMIs depending on whether or not the task pairs were cognate in cyclic environments (i.e., tasks rewarded together or punished together). This allowed us to quantify the magnitude of information about the environment encoded in the mutational neighborhood. Figure 2d shows the total NPMIs for the dominant genotypes from 20 replicates in each regime. The median total NPMIs for *Cyclic* and *Cyclic (Slow)* genotypes were considerably larger than NPMIs from the static regimes (Median for *Const-A*=3.43, median for *Cyclic*=10.28; Mann-Whitney *U* = 46, *p* = 8.17 *×* 10*^−^*^6^, two-tailed). Further, to exclude the influence of perfectly adapted mutants, we recalculated the total NPMIs after removing all perfectly adapted mutants (Fig. S3). Even without the contributions from these mutants, the *Cyclic* and *Cyclic (Slow)* genotypes had larger total NPMIs (Median NPMI for *Const-A*=−0.43, *Cyclic* median=3.59; Mann-Whitney *U* = 52, *p* = 2.138 *×* 10*^−^*^5^, two-tailed). Therefore, evolution under these intermediate-rate predictably-switching environments had a two-fold effect: genotypes were able to create more mutants with an alternate phenotype *and* their local mutational neighborhoods reflected combinations of traits in a manner that mimicked the environmental states the populations encountered over their evolutionary history.

### Evolution along lineages

These results demonstrated that the properties of the mutational neighborhood can dramatically shift during evolution in changing environments. To investigate the implications of these shifts on adapting lineages, we first isolated the lineage of the dominant genotypes, tracking mutation events back to the ancestral genotype. We then calculated the mismatch between the environment state and the phenotypic state of the lineage over evolutionary time, as detailed in the Methods section. Figure 3a shows a moving window average of the mismatch over time. We found that lineages from *Cyclic* and *Cyclic (Slow)* regimes had a lower overall mismatch compared to *Random* and *Cyclic (Fast)* regimes (Median mismatch index for *Cyclic*=0.238, median for *Random*=0.930; Mann-Whitney *U* = 2, *n* = 20, *p* = 5.8 *×* 10*^−^*^11^, two-tailed). Moreover, *Cyclic* lineages show a mismatch only about three times that of *Cyclic (Slow)* even though these lineages evolved in an environment that switched ten times as frequently (Median mismatch index for *Cyclic*=0.238, median for *Cyclic (Slow)*=0.072; Mann-Whitney *U* = 353, *n* = 20, *p* = 9.65 *×* 10*^−^*^6^, two-tailed). To estimate the rate of adaptation, we analyzed the number of elapsed updates before dominant lineages were adapted to an altered environment—a measure we denote as *lineage lag* (see Methods; Fig S1c). In comparison to *Random* or *Cyclic (Slow)*, lineages that evolved under the *Cyclic* regime exhibited a substantially faster phenotypic response following environmental change (median lineage lag for *Cyclic*=3.0, median for *Cyclic (Slow)*=19.0; Mann-Whitney *U* = 50, *p* = 4.94 *×* 10*^−^*^5^, two-tailed; Fig 3b).

**Figure 3:**
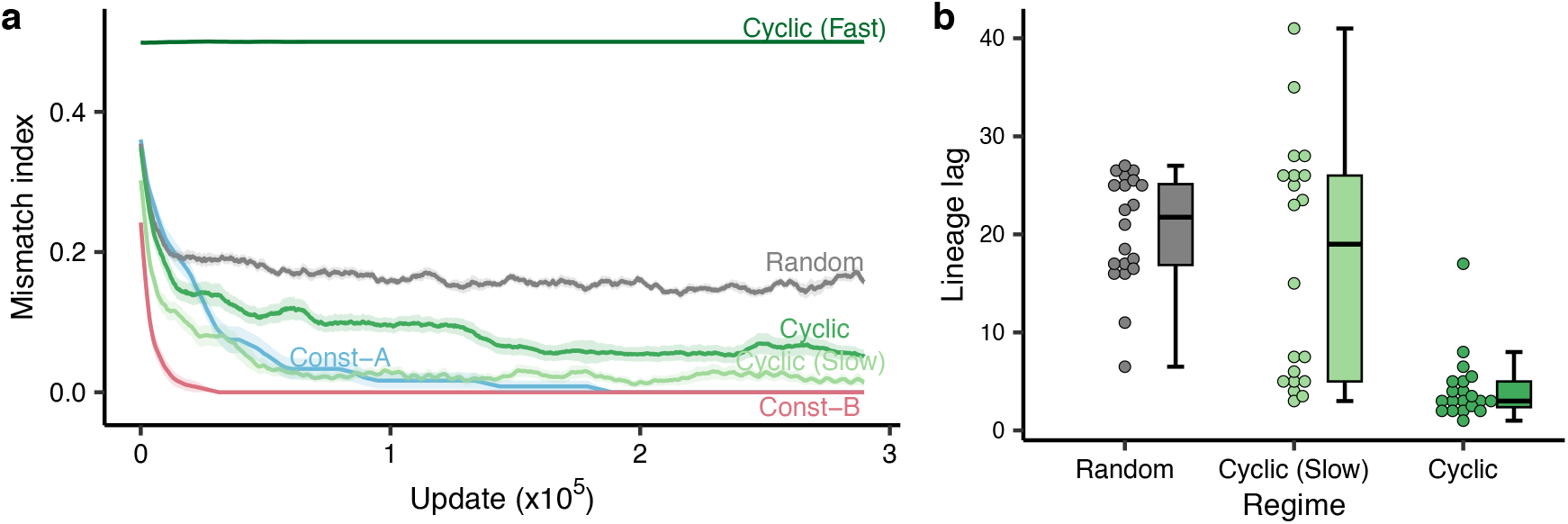
Lineages evolving under specific regimes adapt better and faster to their environments. (**a**) Moving window average of the mismatch between the environment and the lineage phenotypes evolving in different regimes over evolutionary time (window size=10,000 updates). The mismatch index measures the number of task losses or gains required to be perfectly adapted to the lineage genotype’s contemporaneous environment. The central dark line in each curve denotes the mean across replicates, and the width of the ribbons signifies standard error calculated from 20 replicates. (**b**) The lineage lag between the environment and the lineage phenotypes. Lineage lag is calculated as the number of updates elapsed until the lineage achieves a perfectly adapted phenotype after the environment switches. Only lineages from *Random, Cyclic (Slow),* and *Cyclic* regimes are shown here as they consistently achieved perfect adaptation within 100 updates after a switch. Each point is the median of lineage lags calculated from different environmental switches over a single replicate experiment. Box-plot characteristics are the same as in Fig. 2.

We extended our analyses to the mutational neighborhoods surrounding lineage sequences near en-vironmental switch events. An abridged version of this data is presented in figure 4a, with complete data in figure S4. We found that mutational neighborhoods for *Cyclic* lineages appeared to anticipate the environment change; these lineages showed a high concentration of alternate mutants before the environment switched, unlike lineages under the *Cyclic (Slow)* regime that only developed such adaptations gradually after the switch. Synthesizing our observations thus far, we hypothesized that the lineages evolved in cyclic environments localized on phenotypic boundaries in the genotype space, allowing them to switch rapidly between alternate phenotypes (Fig. S1d). Prior theoretical work posits that lineages positioning on such boundaries are able to out-compete lineages farther away from the boundary and thus maximize their long-term fitness [15, 39, 40]. We tested evolved lineages for such localization by calculating the *mutational trade-offs* between adjacent sequences along a line of descent. Essentially, we measured the change in the number of mutants adapted to both environments A and B whenever there was a change in the genotype while moving along the lineage (detailed in Methods). Lineages that localize on boundaries between phenotypes will show the development of strong mutant phenotype trade-offs between adjacent genotypes along their evolved lineages (Fig. S1d). Figure 4b shows these trade-offs as recorded during the last 10,000 generations. Lineages from both *Cyclic* and *Cyclic (Slow)* regimes exhibit trade-offs, with *Cyclic* lineages having the strongest. These trade-offs gradually developed as populations evolved and remained stable for long periods of time (Fig. S5).

**Figure 4:**
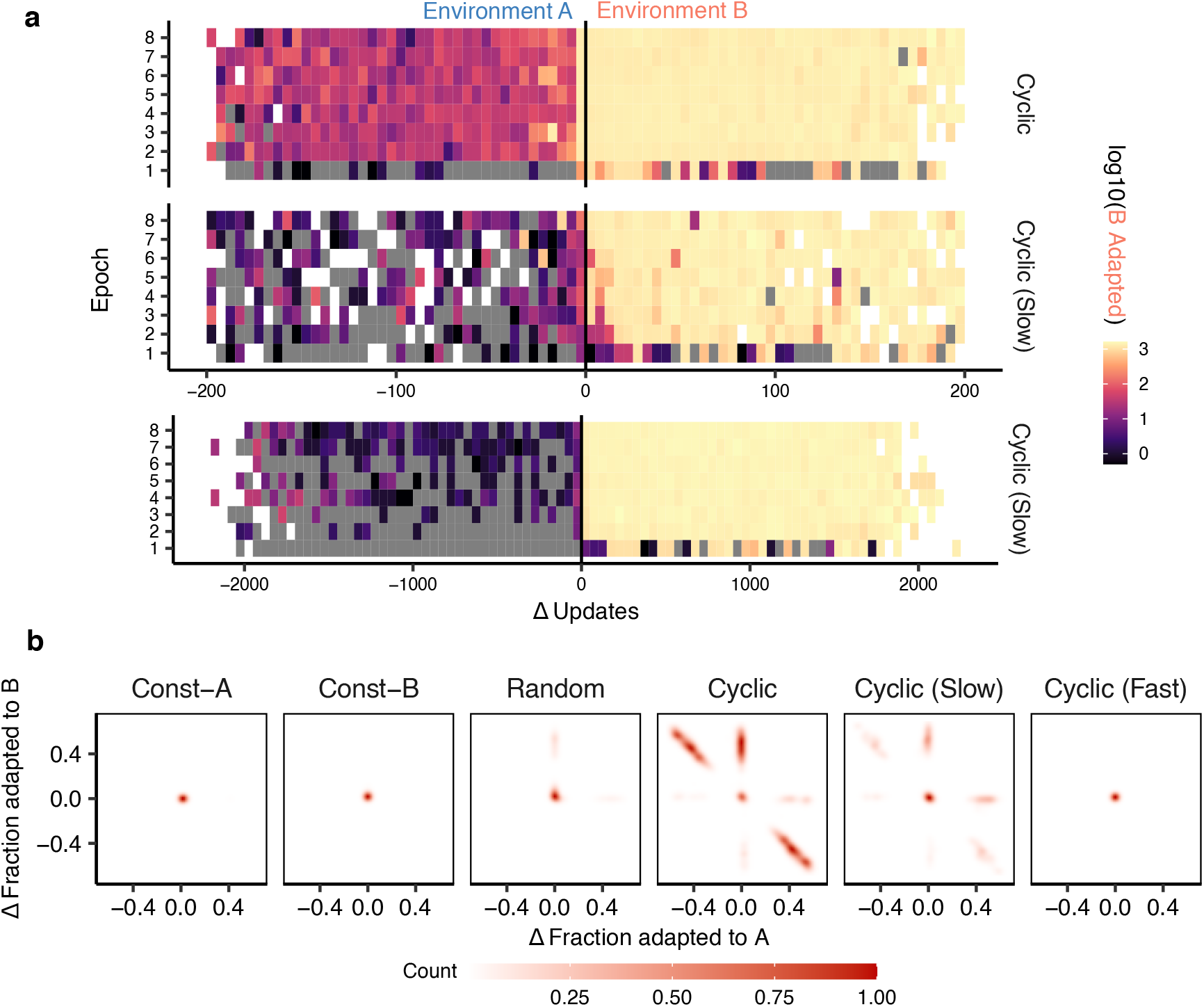
Lineages from the *Cyclic* regime migrate to phenotypic boundaries in the genotype space. (**a**) Median counts of lineage sequence mutants that are adapted to environment B before and after the environment switches from state A to B. The x-axis denotes the number of updates before and after a switch where lineage mutants are recorded. In the top two graphs, each square covers a range of 5 updates. The y-axis segments the entire evolutionary time into eight equal spans, or ‘epochs’. Only *Cyclic* and *Cyclic (Slow)* regimes are plotted here. The lower graph extends the analysis to include up to 2000 updates around the environmental switch for the *Cyclic (Slow)* regime, with the range of each square extended to 50 updates. Grey squares denote points where lineages mutate but do not have any B-adapted mutants. White squares denote points where the lineage does not mutate within the time span of observation. (**b**) Trade-offs between the number of mutants adapted to environment A and B across different regimes during mutation events on the lineage. Points along the *y* = *−x* line denote a 1-to-1 trade-off between mutant phenotype distributions whenever the lineage acquires a new mutation (See Fig. S1d). This plot includes only mutational trade-off data from the final 100,000 updates of evolution. Points from all 20 replicates in each regime are aggregated to calculate the density.

Together, these results demonstrate that selection under changing conditions actively shapes the mutational landscape of populations in a manner that is dependent on the rate and character of environmental change. Both the accessibility of alternative phenotypes and the information stored in the distribution of mutant phenotypes are substantially enhanced during this process. These features, in turn, help the population adapt quickly when facing ongoing environmental fluctuations.

### Evolution of mutation rates

Mutation rates in natural populations are known to evolve rapidly under stressful conditions [41, 42]. However, the drift barrier hypothesis proposes that the mutation rate should decrease to a minimal value sustained above zero by genetic drift [22]. To explore the dynamic interaction between mutation rate evolution and the evolution of the mutational neighborhood, we conducted a series of experiments allowing the mutation rates to evolve naturally. In Avida, organisms inherit their parent’s mutation rate, subject to small stochastic variation proportional to their parent’s rate (see Methods, Fig. S1e). To mitigate bias introduced by any specific starting mutation rate, we initiated experiments with rates spanning several orders of magnitude (0.0001, 0.001, and 0.0316). Then, we measured the average mutation rate of organisms in these populations as they evolved under different regimes. Figure 5a shows the average evolved mutation rates for populations starting from three different initial values. Environments undergoing intermediate fluctuations—specifically, the *Cyclic* and *Random* regimes— were conducive to the evolution of higher mutation rates (with initial mutation rate of 10*^−^*^3^, median evolved mutation rate in *Cyclic*=9.19*×*10*^−^*^4^, median in *Const-A*=7.37*×*10*^−^*^6^; Mann-Whitney *U* = 400, *n* = 20, *p* = 1.45 *×* 10*^−^*^11^, two-tailed). This trend toward greater mutation rates in *Cyclic* held true across all starting mutation rates studied (with an initial mutation rate of 10*^−^*^4^, median evolved mutation rate for *Cyclic*=1.88 *×* 10*^−^*^4^, median for *Const-A*=1.611 *×* 10*^−^*^5^; Mann-Whitney *U* = 387, *n* = 20, *p* = 5.41 *×* 10*^−^*^9^, two-tailed).

**Figure 5:**
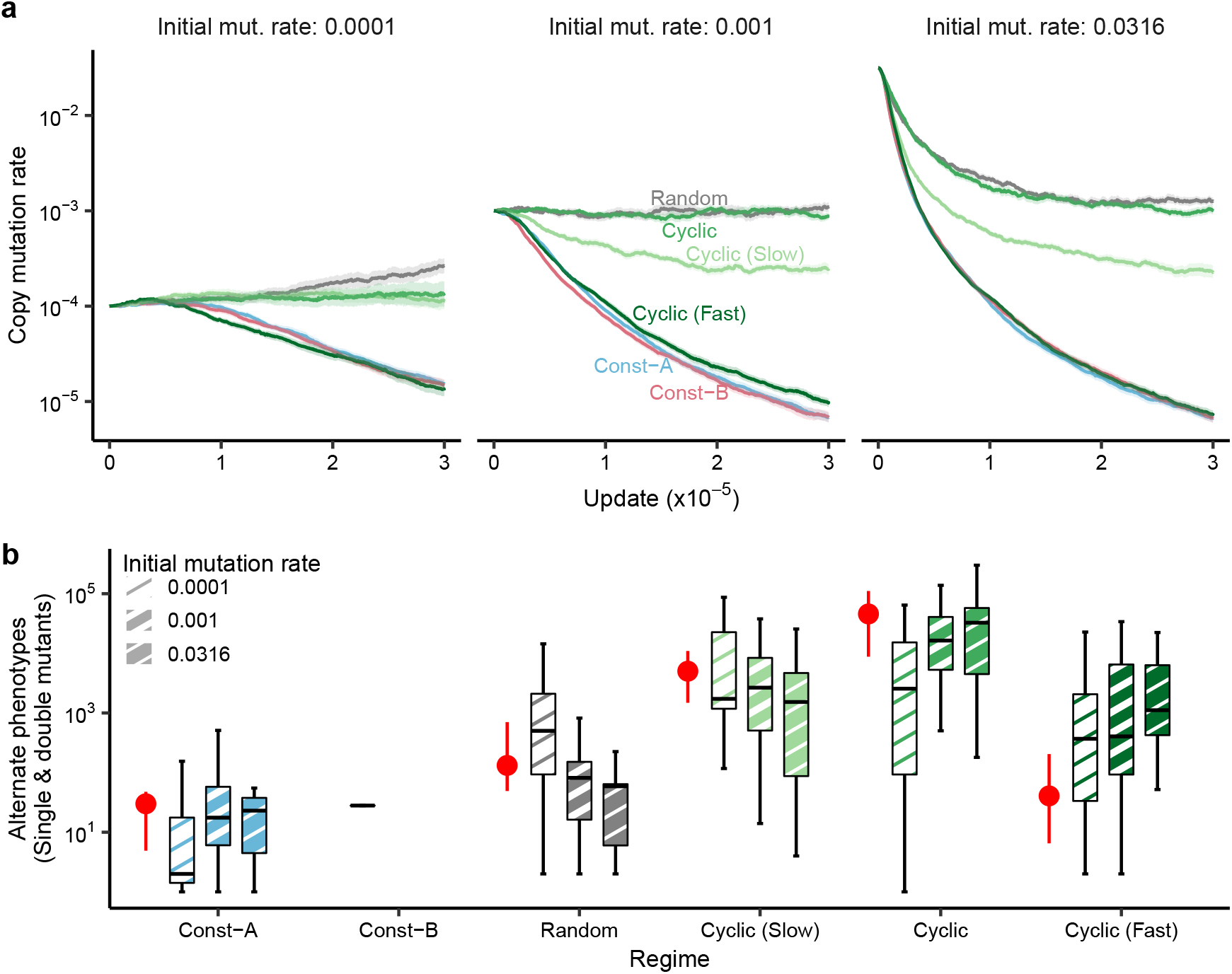
Concurrent evolution of high mutation rates and accessible mutational neighborhoods in fluctuating environments. (**a**) Mean mutation rate for populations evolved under different regimes of environmental change. Dark lines represent the average mutation rates calculated from 20 distinct replicates, while the ribbons denote confidence intervals defined by two times the standard error. The three panels show the data from three sets of experiments that were started with different initial mutation rates. (**b**) The total number of single and double mutants of the end-point dominant genotype with an alternative phenotype. The patterned box plots categorize dominant genotypes according to the initial mutation rates used in the experiments. Other box plot characteristics are the same as in Fig. 2. The red filled circles denote the median number of alternate phenotype mutants from the fixed mutation experiments, with the error bars denoting the lower and upper quartiles (allowing direct comparison to the data from Fig. 2b).

Elevated mutation rates can increase the mutational load in populations by raising the incidence of deleterious genotypes, thereby diminishing the average fitness within a static environment [43]. Hence, we anticipated a detrimental impact of high mutation rates on the accessibility of beneficial alternative phenotypes. Counter to this expectation, the *Cyclic* regime populations maintained increased access to mutants adapted to the alternative environments, while simultaneously elevating their mutation rates (Fig. 5b; for initial mutation rate of 10*^−^*^3^, median number of alternate mutants for *Cyclic*=16378.5, median for *Const-A*=0.0, *Const-A* mean=27.5; Mann-Whitney *U* = 399, *n* = 20, *p* = 2.86 *×* 10*^−^*^8^, two-tailed). The number of alternate mutants in the evolving mutation rate experiments were comparable to the fixed mutation rate case (median number of alternate mutants in *Cyclic* with fixed mutation rate=45802.5, *Cyclic* with an initial mutation rate of 0.001 median=16378.5; Mann-Whitney *U* = 139, *n* = 20, *p* = 0.102, two-tailed). The evolved NPMIs were also similar to genotypes from the fixed rate treatment (Fig. S6). Thus, populations evolved higher mutation rates and greater accessibility of adaptive variation for alternative environments simultaneously in the *Cyclic* regime.

### Adaptive performance under alternative fluctuations

Populations typically excelled when subjected to the same environmental conditions that shaped them; however, it remains unclear if such specific adaptations confer benefits across varying ecological conditions. To ascertain whether adaptation within a certain regime delivered cross-regime fitness improvements, we isolated genotypes from the initial evolution experiments and subjected them to each alternative regime. Due to computational constraints, the evolving mutation rate treatment was represented by genotypes from the intermediate initial mutation rate treatment only, with a starting mutation rate of 0.001. To measure the evolutionary performance of genotypes, we calculated the average mismatch index and the average lineage lag over approximately 1,000 generations of evolution. The data is summarized in figure S7.

We found that genotypes evolved in the *Cyclic* regime had a lower average mismatch and adapted faster following environmental changes than their counterparts from the *Cyclic (Slow)* or *Cyclic* regimes (Fig. S7). Contrary to expectations, genotypes from variable mutation rate experiments perform worse those from the fixed mutation rate experiments. This decrease in performance may be attributable to the slight reduction in the evolved mutation rates compared to the fixed rates (basal mutation rate=10*^−^*^3^, median evolved mutation rate in *Cyclic* with intermediate starting mutation rate=9.19 *×* 10*^−^*^4^). Nevertheless, the possibility remained that adaptation via mutation rates may come at the cost of evolvability via the mutational neighborhood — a hypothesis we examine further in the concluding section of our results.

### Adaptation in novel environments

To measure how the ability of populations to adapt to new environments changed during our evolution experiments, we subjected evolved genotypes to a novel environment with 127 unique rewarded tasks unrelated to those rewarded in environments A or B. During 30,000 generations of evolution, we measured the total number of tasks performed by the descendants of evolved genotypes, initially adapted to different environmental regimes. Data reflecting the progression of task acquisition over time is shown in S7c. We found that no marked differences in the evolvability of genotypes from the fixed mutation rate experiments to this new environment, as indicated by the number of new tasks they perform (Kruskal-Wallis *χ*^2^ on the number of tasks evolved at the end, for fixed mutation rate genotypes=4.1292, *df* = 5, *p* = 0.531).

By contrast, genotypes from the evolving mutation rate experiments exhibited significant diversity in their adaptive success in this new environment (Kruskal-Wallis *χ*^2^ for the number of tasks evolved for evolving mutation rate genotypes=94.221, *df* = 5, *p <* 2.2 *×* 10*^−^*^16^). Notably, genotypes from *Random* and *Cyclic* regimes that underwent mutation rate escalation acquired more functions than their ancestral genotypes (median tasks evolved by ancestral genotype=30, median tasks evolved by *Cyclic* genotypes = 42.3, 18 of 18 samples evolved more tasks than the ancestral median, *p* = 7.629 *×* 10*^−^*^6^, two-tailed one-sample sign test). Inversely, genotypes conditioned to *Const-A*, *Const-B*, and *Cyclic (Fast)* evolved fewer tasks compared to their ancestor (median tasks evolved by *Cyclic (Fast)* genotypes = 17.9, 19 of 19 samples evolved fewer tasks than the ancestral median, *p* = 3.815 *×* 10*^−^*^6^, two-tailed one-sample sign test).

To test whether mutation rate was the key driver of the observed differences, we transplanted the evolved mutation rates into organisms from the fixed rate experiments. This exchange recapitulated the difference in the rate of adaptation seen between treatments in this new environment (Fig. S7c; Kruskal-Wallis *χ*^2^ for the number of tasks evolved at the end for fixed mutation rate genotypes transplanted with evolved mutation rates=95.167, *df* = 5, *p <* 2.2 *×* 10*^−^*^16^). Conversely, when the evolving mutation rate genotypes were transplated with the ancestral mutation rate, their rate of adaptation remained largely unchanged (Kruskal-Wallis *χ*^2^ for the number of tasks evolved at the end for the evolution mutation rate genotypes transplanted with the ancestral mutation rates=3.5192, *df* = 5, *p* = 0.6205). In conclusion, the data suggests that while the ability to access alternate phenotypes provides a mechanism to quickly adapt to previously encountered environments, elevated mutation rates are a primary driver of adaptation to entirely new environments.

### Interaction between evolution of the mutation rate and the mutational neighborhood

The mutation rate is a key determinant of the rate at which populations explore genetic landscapes and is a regulator of their equilibrium genetic diversity [44, 45]. Thus, we expect that different mutation rates would lead to distinct evolved mutational neighborhoods. To understand the role of the mutation rate in the evolution of mutational neighborhoods, we evolved twenty replicate populations at five different fixed mutation rates in three different environmental regimes. One selected mutation rate mirrored the evolved mutation rate from previous experiments (*µ* = 9.19*×*10*^−^*^4^). The number of alternate phenotype mutants that evolved in different regimes under different values of the mutation rate are shown in figure 6. We found that the effect of the mutation rate on the evolution of the mutational neighborhood depends on the way the environments changed. In the static *Const-A* regime, only a small number of the populations evolved alternate phenotypic mutations. In the *Cyclic (Slow)* regime, increasing the mutation rate decreased the abundance of alternate phenotypic mutations (*p* = 0.00104, see Table S1). This observation is perhaps expected since frequent mutations would favor populations in more robust regions of the genotype-phenotype map (e.g., “survival of the flattest”) [46]. In these gradually shifting environments, the drive towards robustness seemed to impede the evolution of the mutational neighborhood as mutation rates increased.

**Figure 6:**
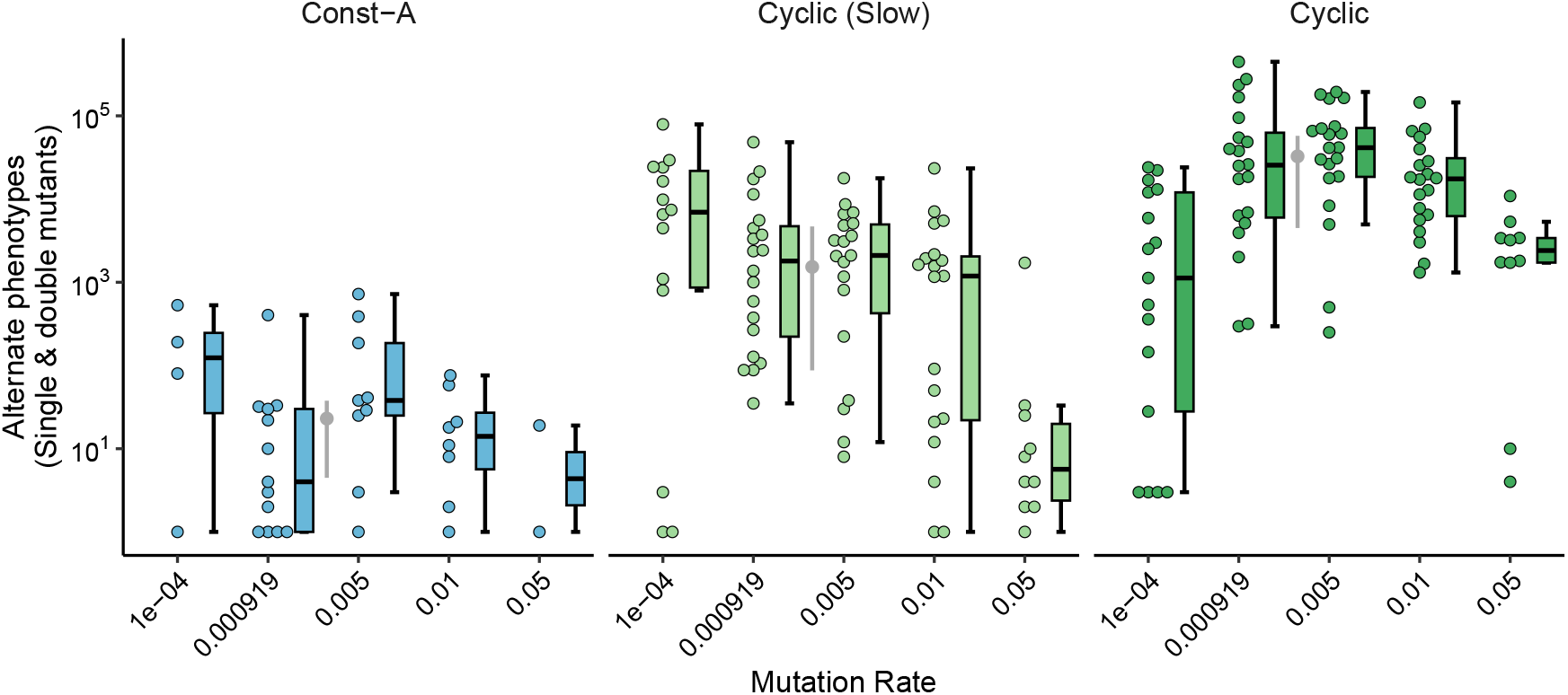
Interplay of mutation rate and the mode of environmental change shapes the mutational neighborhood. This graph plots total number of single and double mutants of the end-point dominant genotype with an alternative phenotype. The data is plotted for evolution experiments in three different regimes with different values of a fixed mutation rate (x-axis). The mutation rate of 9.19×10^−4^ was set to mimic the evolved mutation rate in previous experiments (specifically, in the *Cyclic* regime starting with an initial mutation rate of 10^−3^). The filled grey circle and error bars denote the mean and quartiles of alternate phenotype counts observed during the evolving mutation rate experiments.

Surprisingly, we found a non-monotonic relationship between mutation rate and the number of alternate phenotypic mutants in the *Cyclic* regime (*p* = 2.39×10^−05^, see Table S1). The number of alternate phenotypic mutants first increased with mutation rate but subsequently decreased at the highest mutation rates, with an estimated peak at a mutation rate of 0.0018. Notably, we see the largest increase in the number of alternate phenotypic mutants at a similar mutation rate to which the populations evolved in our evolving mutation rate experiments. On one hand, the negative correlation between alternate phenotypes and mutation rate at high values of the mutation rate mirrors the results in more slowly varying environments. On the other hand, the number of alternate phenotypes increased over a large range of mutation rates in this environment. Considering the number of alternate mutants in *Cyclic* is always higher than the *Slow* regime, these results suggest synergy rather than antagonism between the evolution of mutation rates and the availability of alternate phenotype mutations in environments fluctuating at an appropriate speed.

## Discussion

Here, we experimentally study the evolution of the primary determinants of evolvability simultaneously (the *mutational neighborhood* and the *mutation rate*), uncover their interaction with one another, and tie their emergence to the mode of environmental change. We show that evolved organisms can reside in mutational neighborhoods with an elevated number of alternate phenotypes that allow them to adapt to changing environments quickly. This occurs when evolving populations move to areas of the genotype space where such phenotype-altering mutations are more abundant—i.e., to *phenotype boundaries* (see Fig. 1c for an illustration). Importantly, this shift does not occur from directly biasing mutations towards particular phenotypes as modifier theory posits [47]. Rather, the phenotypic bias we observe manifests indirectly from structural changes in the mutational neighborhood. The movement towards phenotypic boundaries is most readily evolved in environments that switch between *predictable* states at *intermediate* rates. By predictable, we mean that future environments reliably resemble those encountered previously. In our experiments, the ideal rate for evolvability is realized when the environment switches between two states approximately every 30 generations. Therefore, our results indicate that evolution in certain environments can favor genotypes that are not necessarily more fit presently, but rather have a propensity to generate more adaptive mutants in the future.

We devise a mutual information metric that captures the extent to which the mutational neighborhood reflects the mode of environmental change. Using this metric, we find that the distribution of mutant phenotypes becomes tuned to environments encountered by the population during evolution. Thus, we show that the set of phenotypes available to a population via mutations evolves to store information about environmental change, allowing the population to “learn” how to adapt to recurring environments [48]. We note an interesting macroevolutionary consequence of this result, where one would expect a lineage that had adapted to changing environments to show greater phenotypic variation than lineages adapted to more stable environments (Fig. S9 presents brief evidence of this in our system). As a corollary, the variation revealed through mutation can offer clues to the kinds of environments encountered by lineages in the past [33, 49]. These results provide a link between the microevolutionary processes that generate adaptive mutations and patterns that manifest at macroevolutionary scales.

Elevated mutation rates can limit adaptation by increasing the accumulation of deleterious mutations [43]. Yet, the frequency of such deleterious mutations is also contingent upon an organism’s local mutational neighborhood. In agreement with the drift barrier hypothesis, we find that the mutation rate precipitously decreases to minuscule values in static environments or environments changing very rapidly [22]. Remarkably, however, we see that the populations maintain much higher mutation rates in environments changing at intermediate rates. In these environments, the evolved mutation rates approximated those required for the evolution of the most evolvable mutational neighborhoods (as inferred from static mutation rate experiments). In contrast to previous work in which evolution failed to optimize mutation rates in a static environment, we show that such optimization is possible in more complex fluctuating environments [50]. The effect of fluctuations on evolving mutation rates was also observed in a recent study, where populations evolved elevated mutation rates when population bottlenecks were applied at intermediate frequencies during experimental evolution [23]. Whether the observed mutation rate increase in our study derives from a similar mechanism — specifically, the pruning of maladaptive phenotypes during environmental shifts — remains an open question.

Counter to our expectations based on their compounding effects on mutational load, we observe increases in both the mutation rate and the accessibility of alternate phenotypic mutants in the recurrent and intermediately switching environments[18]. The benefits of these evolvability mechanisms manifest in different ways. On one hand, evolved accessibility of alternate phenotypes helps populations rapidly adapt to previously encountered environments, thus exhibiting a “memory-like” response to changes. Furthermore, organisms with such evolvability demonstrate quicker adaptation across a range of environmental change rates, not just the rate they historically experienced. Elevated mutation rates allow quick adaptation in completely new environments. Furthermore, in these novel environments, organisms starting with the same mutation rate but different mutational neighborhoods ultimately achieve similar fitness. When the mutation rate is artificially forced high, mutational neighborhoods evolve few alternative phenotypes, except when the environment fluctuates at intermediate rates. Thus, intermediate-rate environmental fluctuations are able to overcome the otherwise overwhelming selection for mutational robustness. Together, these mechanisms of evolvability prime genotypes for rapid adaptation to both historical and novel environments.

The emergence of evolvability is inherently tied to the structure of genotype space [51–54]. Boundaries between distinct regions in genotype space that encode different phenotypes are central to the evolvability that we study, but their presence in every genotype-phenotype map is not guaranteed [55–58]. Moreover, such boundaries may not always be accessible through evolution from the ancestral genotype. It is essential to address the abundance of such structural features in the genotype-phenotype map for more general conclusions to be drawn [35, 59]. Previously studied models demonstrate that intermediate rates of environmental change influence population localization, but these results are derived from necessarily simple models under idealized assumptions [15, 39]. Our work demonstrates that this effect can be readily observed in a realistic–but quite unnatural–evolving system with a markedly complex genotype-phenotype map and identifies the kinds of environmental change regimes that favor such localization.

In conclusion, we show that evolution under specific modes of environmental change promotes high mutation rates and simultaneously shapes the local mutational neighborhood of populations. By measuring the rate of adaptation of evolved organisms in new environments, we show that increases in evolvability acquired through these two mechanisms play distinct roles in historical and novel environments. Using large-scale digital evolution experiments, we trace the dynamics and interplay of these modes of evolvability as they evolve, and show that this escalation is characterized by the movement of populations towards boundaries in genetic space. In subsequent work, we aim to analytically resolve the dependence of evolvability on the environment, and test if our results can be harnessed for the directed evolution of populations with greater evolvability. Although we use an unnatural study system, our results provide insight into how and why evolution in nature can evolve to be so relentless.

## Methods

### *In silico* experiments

We used the Avida Digital Evolution Platform to perform *in silico* evolution experiments. Avida organisms are autonomous, asexual, self-replicating computer programs that compete for space and execution time in a virtual environment [34]. These organisms possess a genome—a linear sequence of computer instructions—that enables them to reproduce and perform computations (Fig. S1a). The Avida instruction set includes operations for basic computations, flow control (e.g., conditional logic and looping), input, output, and the building blocks of self-replication. To reproduce, organisms must copy their genome instruction-by-instruction and then divide. Copy operations are imperfect and result in single-instruction mutations in the offspring genome. Time in Avida is measured in *updates*. On average, an update spans the execution of 30 instructions in an organism’s genome. After division, new offspring are placed randomly in the world, replacing any existing occupants. Organisms thus compete for space, which drives selection to improve their replication rate in order to not be overwritten by competitors. Performing computations allows organisms to acquire a fitness multiplier depending on the environment.

The average generation time of an Avida organism in our experiments is around 10 updates. We ran the primary evolution experiments in this work for a total of 300,000 updates (around 30,000 generations) with a carrying capacity of 22,500 organisms. The two environments in our experiments— A and B—rewarded an alternate set of computations (*tasks*) (Fig. S1b). Environment A rewarded the organisms with a fitness multiplier of 2.0 if they performed tasks NOT, AND, or OR - and penalized NAND, ORN, or ANDN with a multiplier of 0.8. Environment B rewarded the organisms with a fitness multiplier of 2.0 for NAND, ORN, or ANDN - and penalized NOT, AND, or OR with a 0.8 multiplier. The rewards and penalties stacked, thus providing the maximum fitness benefits to organisms that were perfectly adapted to perform only the tasks rewarded in each environment.

We initiated all primary evolution experiments with identical ancestral organisms having a genome that was 100 instructions in length, which was fixed at this size. For experiments under a fixed mutation rate, we used a copy mutation rate of 0.001, which created approximately one substitution out of every 1000 instructions copied. For experiments where mutation rates were allowed to evolve, we started with three different initial mutation rates—0.0001, 0.001, and 0.0316. New offspring inherited the mutation rate of their parents, but were modified by adding a deviation drawn from a Gaussian distribution with a mean of 0 and a standard deviation of 1% of the parent’s mutation rate value (Fig. S1f) [50].

### Environment change regimes

The environment change regime determines the way environments switch in an experiment (Fig. S1b). The *Const-A* and *Const-B* regimes were in environmental states A and B, respectively, throughout evolutionary time. The *Cyclic* regime switched between environments A and B, with the waiting time between switches picked from a Gaussian distribution with a mean of 300 updates (*∼*30 generations) and a standard deviation of 60 updates. The *Cyclic (Fast)* regime switched between A and B with a mean waiting time of 30 updates (*∼*3 generations) and a standard deviation of 6 updates. The *Cyclic (Slow)* regime switched between A and B with a mean waiting time of 3000 updates (*∼*300 generations) and a standard deviation of 600 updates. In the *Random* regime, tasks to be rewarded or penalized were redrawn randomly from the set of six tasks described before, with a mean waiting time of 300 updates (standard deviation of 60 updates). All regimes ended with 300 updates of evolution in a constant terminal environment, which was environment B for Const-B, and environment A otherwise. Evolution in these terminal environments allowed us to compare adaptive states at the end of the experiments consistently across regimes.

### Characterizing the mutational neighborhood

We performed an exhaustive survey of the 2-step mutational neighborhood of the dominant genotype from the end of the experiments (Fig. S1c). To do so, we created every possible single- and double-point mutation in the dominant genotypes isolated after evolution. We tested each mutant for viability (i.e., the ability to self-replicate) and measured the phenotypes for viable mutants (i.e., the set of tasks the mutant could perform). We also isolated the lineages of the dominant genotypes and performed a similar mutational neighborhood analysis on the entire lineage. Only single-point mutations were assayed for lineages. The mutant phenotype data from endpoint dominant genotypes was used to calculate the *pointwise mutual information*. The mutant phenotype data from lineages was used to measure the *mismatch index*, *lineage lag*, and *mutational tradeoffs* (see below).

### Pointwise mutual information

The pointwise mutual information (PMI) for two events *x, y ∈* Ω is defined as,

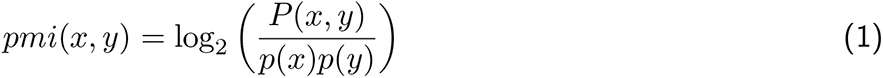

Where Ω is the sample space of events, *p* is the probability mass function on Ω, and *P* is the probability mass function on the joint space Ω × Ω. We normalized this value between −1 and 1 by defining the normalized pointwise mutual information (NPMI) as,

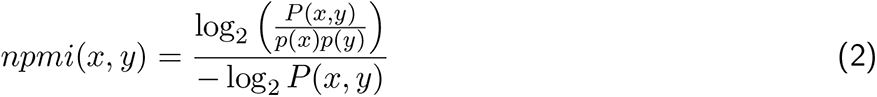

The NPMI measures the excess probability of occurrence for certain event pairs in the joint probability distribution when compared to the expected probabilities based on these events’ marginal distribution. For the measurements in this work, *x* and *y* denote the different tasks in the set of possible tasks recognized by the environments. *p*(*i*) was measured as the fraction of all possible double mutants that performed the task *i*. *P* (*i, j*) was measured as the fraction of all possible double mutants that performed both tasks *i* and *j*. A positive NPMI indicates that a task pair is over-represented in the mutants than expected based on the occurrence of each individual task. A negative NPMI indicates that the task pair is underrepresented on average.

The joint sample space *S* = Ω × Ω consists of 15 unique, non-similar, task pairs. These pairs are labeled as cognate (*∈ S_C_*) or non-cognate (*∈ S_N_*) depending on whether they were rewarded by the same environment or not, respectively. The total NPMI was calculated as the sum of individual NPMIs after assigning the task pair a sign based on its cognate/non-cognate label.

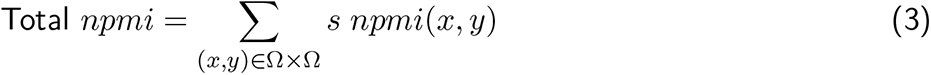

Where,

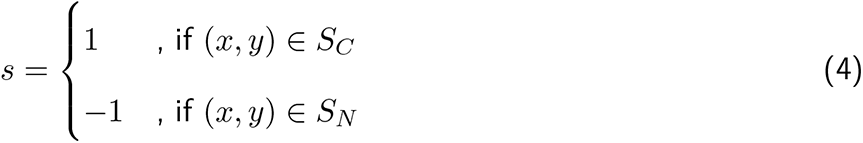

We use this total NPMI metric to compare the information encoded on the mutational neighborhood of the dominant genotypes obtained from different regimes.

### Lineage measurements

We calculated the mismatch index for lineages as the Hamming distance between the lineage phenotype and the phenotype most fit in the current environment. Hamming distance between phenotypes measured the minimum number of task losses/gains required to change between two phenotypes. For example, an organism that performs NOT, AND, and OR is three units away from an environment that has the maximum fitness for organisms performing NOT and NAND (two losses and one gain of function required). We further averaged the mismatch index over multiple updates to smooth out jumps in the index that happened due to sudden environmental changes.

The lineage lag was calculated to assay the time required for a lineage to adapt to a changed environment (Fig. S1d). We surveyed 100 updates both before and after environment switches to see if the lineage ever acquired a phenotype perfectly fit for the new environment. Then, we calculated the amount of time between when the lineage became perfectly adapted and the environment switch. This number is positive if the lineage switched after the environment changed and negative if it switched before the environment.

We measured the mutational trade-offs along the lineage by calculating the change in the fraction of mutants adapted to environment A (Δ*A*) and environment B (Δ*B*) between genotypes occurring along the dominant lineage (Fig. S1e). Points with coordinate (Δ*A,* Δ*B*) were then plotted in a 2D Cartesian plane. Points along the axes of the 2D plane indicate mutation events along the lineage where an increase or decrease in mutants with one adaptation was not accompanied by a change in the number of mutants with the other adaptation. A high density along the *y* = *−x* line denotes a 1-to-1 trade-off between mutations, where an increase in mutants adapted to one environment came at a cost of the number of mutants adapted to the other environment (Fig. S1e).

### Cross-regime experiments

We evolved the dominant genotypes—isolated from both fixed and evolving mutation rate experiments— under different regimes to test their performance directly. We released a single organism with the isolated genotype in a new Avida world where the environment switched as described by the four fluctuating regimes - *Cyclic*, *Cyclic (Slow)*, *Cyclic (Fast)*, and *Random*. Mutation rates were set at the start of the experiment based on whether the genotypes were isolated from a fixed or evolving mutation rate experiment but were not allowed to evolve further. These Avida worlds had a carrying capacity of 22,500 organisms (same as the primary evolution experiments) and the experiments were run for 10,000 updates. The performance of these genotypes was measured by calculating the time-averaged mismatch and average lineage lag as described earlier.

### Measuring evolvability in new environments

To test the evolvability of organisms that evolved under different regimes in new environments, we released the dominant genotypes in an environment where 127 novel tasks—distinct from the 6 tasks encountered during primary evolution—were rewarded. We then performed a secondary evolution experiment for 300,000 updates where the organisms evolved under a fixed mutation rate. For genotypes isolated from fixed mutation conditions, we used a basal mutation rate of 0.001. For genotypes isolated from evolving mutation rate conditions, the mutation rate was set to the rate they had evolved in the primary evolution experiments. To account for the direct effect of mutation rates, we also performed *reciprocal* experiments where the organisms that evolved in evolving mutation rates were assigned a mutation rate of 0.001, and the organisms that evolved under fixed mutation rates were assigned the mean mutation rate that evolved in the evolving mutation rate treatments.

### Statistics

Statistical analyses were performed using functions in base R [60]. The correlation plot in figure S2 was generated using the corrplot package. Details about the statistical tests, including the type of test used, test statistic value, and p-values are provided in the text. For more details and the code used to perform statistical analyses, please refer to the “Plotter/Statistics file path” field in the data guide mentioned below.

## Data Availability

The data generated for and used in this work has been archived publicly at https://doi.org/10.17605/OSF.IO/7P9KX. The data guide.pdf file (found in the root directory of the repository) is a data guide that annotates the provided data. It points to scripts used to run experiments, analyze data, perform statistics, and plot the figures. Interested readers can follow the steps mentioned in the data guide and repository to reproduce the data and figures in this paper.

## Code Availability

Avida, the open-source digital evolution framework used in this work, is available under the MIT license at https://github.com/devosoft/avida. All data, scripts, and analysis files used in this work have been archived publicly. They are available in the repository mentioned under Data availability.

## Acknowledgments

We thank Dr. Jordan Horowitz for his valuable comments. This work was supported by the computational resources and services provided by Advanced Research Computing at the University of Michigan, Ann Arbor. This research was supported in part by the National Science Foundation grant DEB 1813069.

## Author Contributions

B.K. and L.Z. wrote the manuscript. B.K., A.L., M.A., and L.Z. conceptualized the project, designed the experiments, and ran the analyses. All authors edited and reviewed the manuscript. L.Z. supervised and funded the work.

## Supplementary

**Figure S1:**
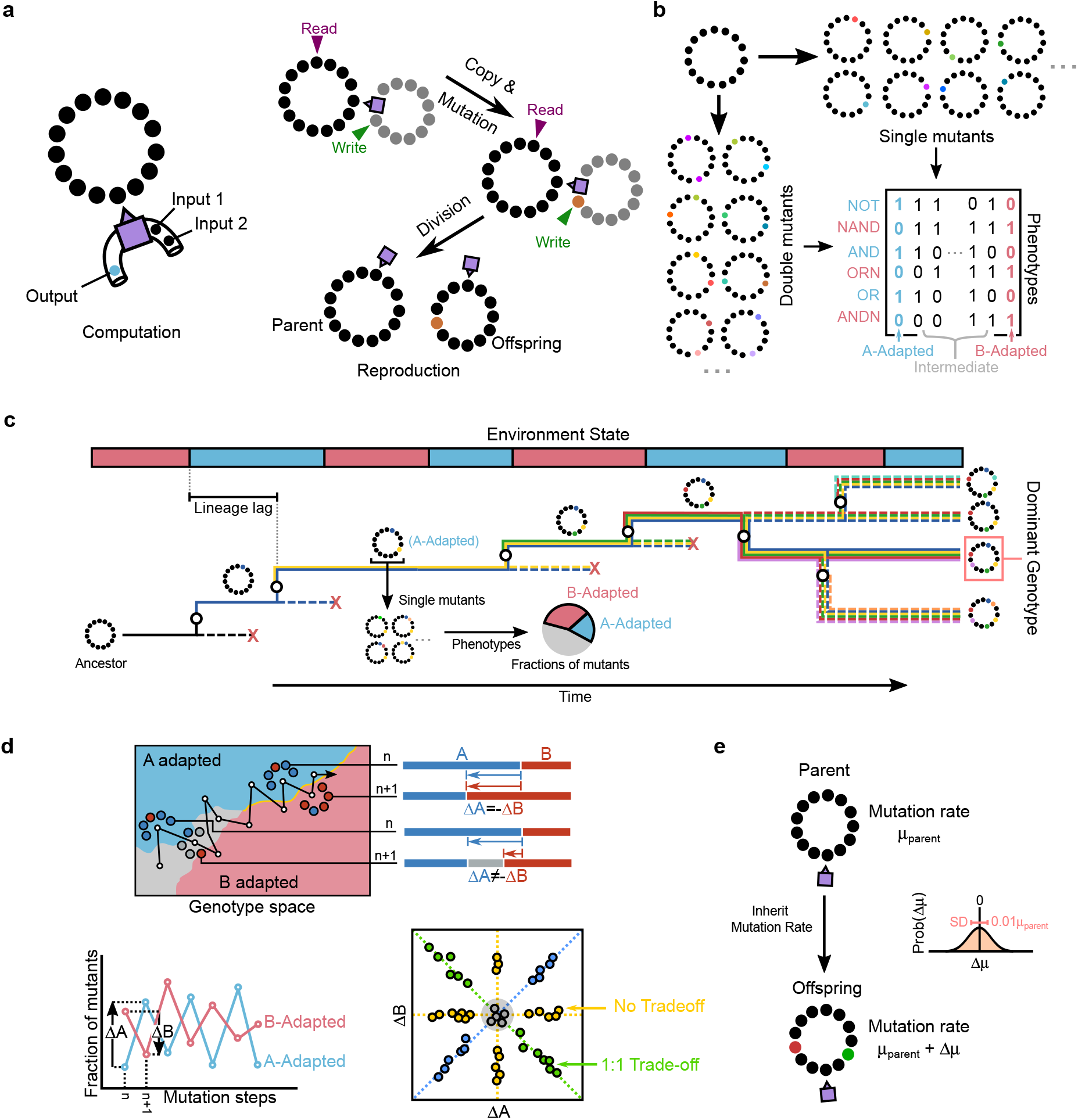
In silico evolution in Avida and experimental methods. (**a**) An Avida world consists of multiple genetic sequences, encoded as a circular series of instructions (black circles) that can be executed by a virtual CPU (purple square) that translates these instructions into organismal function. The main organismal functions include *computation* - where organisms perform logical tasks to increase reproductive efficacy - and *reproduction* - where the sequence instructs the CPU to copy instructions into a new memory location (grey and red circles). During computation, the sequence requests inputs from the environment and acts upon these inputs to generate an output. We use two environments in this study - A and B - both of which recognize a set of six possible computations or tasks. Refer to Fig. 2a for more details about the environment states and regimes. (**b**) To survey the mutational neighborhood of evolved organisms, we create every possible single (top-right) and double mutant (bottom-left) and measure their phenotype. Phenotypes denoting tasks that are rewarded in environments A or B are labeled *A-adapted* or *B-adapted* respectively. All other phenotypes are classified as *intermediate*. Sequences that cannot self-replicate are classified as an *inviable* phenotype (**c**) An example of a pruned phylogeny of digital organisms evolving in a cyclically changing environment. The colors on a branch denote the set of mutations acquired by the branch compared to the ancestral genome. For most of the data, we look at the lineage of the dominant genotype which is shown as solid lines. Circles on the phylogeny denote mutation events where the phenotype of the lineage may or may not change. *Lineage lag* is calculated as the time it takes for the lineage to adopt a perfectly adapted phenotype following an environment switch, as shown on the top left. When calculating mutational neighborhoods for the lineages, we only look at the single mutants of the lineage genotypes. (**d**) A hypothesized mechanism of localization that promotes increased access to alternate phenotypes (top). Lineages start in the intermediate adapted region of a genotype space (grey) but slowly find regions where a few mutations allow quick changes in phenotype (yellow curve). If this hypothesis holds, we expect the development of increased mutational trade-offs between consecutive mutational steps on the lineage—i.e., an increase in lineage mutations where the increase in number of mutants adapted to environment A is accompanied by an equivalent decrease in the number of mutants adapted to environment B. We check for these trade-offs by measuring the number of mutants of lineage sequences adapted to environments A and B (bottom-left) and calculating the differences (Δ*A* and Δ*B*). When plotted on a Cartesian plane, mutation events that show trade-offs appear along the *y* = *−x* line. (**e**) Schematic of the mechanism for evolving mutation rates. Offspring inherit the mutation rate by sampling from a Gaussian distribution with a mean that is equal to the parent’s mutation rate (*µ*_parent_) and a standard deviation that is 1% the parent’s mutation rate (Δ*µ* = 0.01*µ*_parent_).

**Figure S2:**
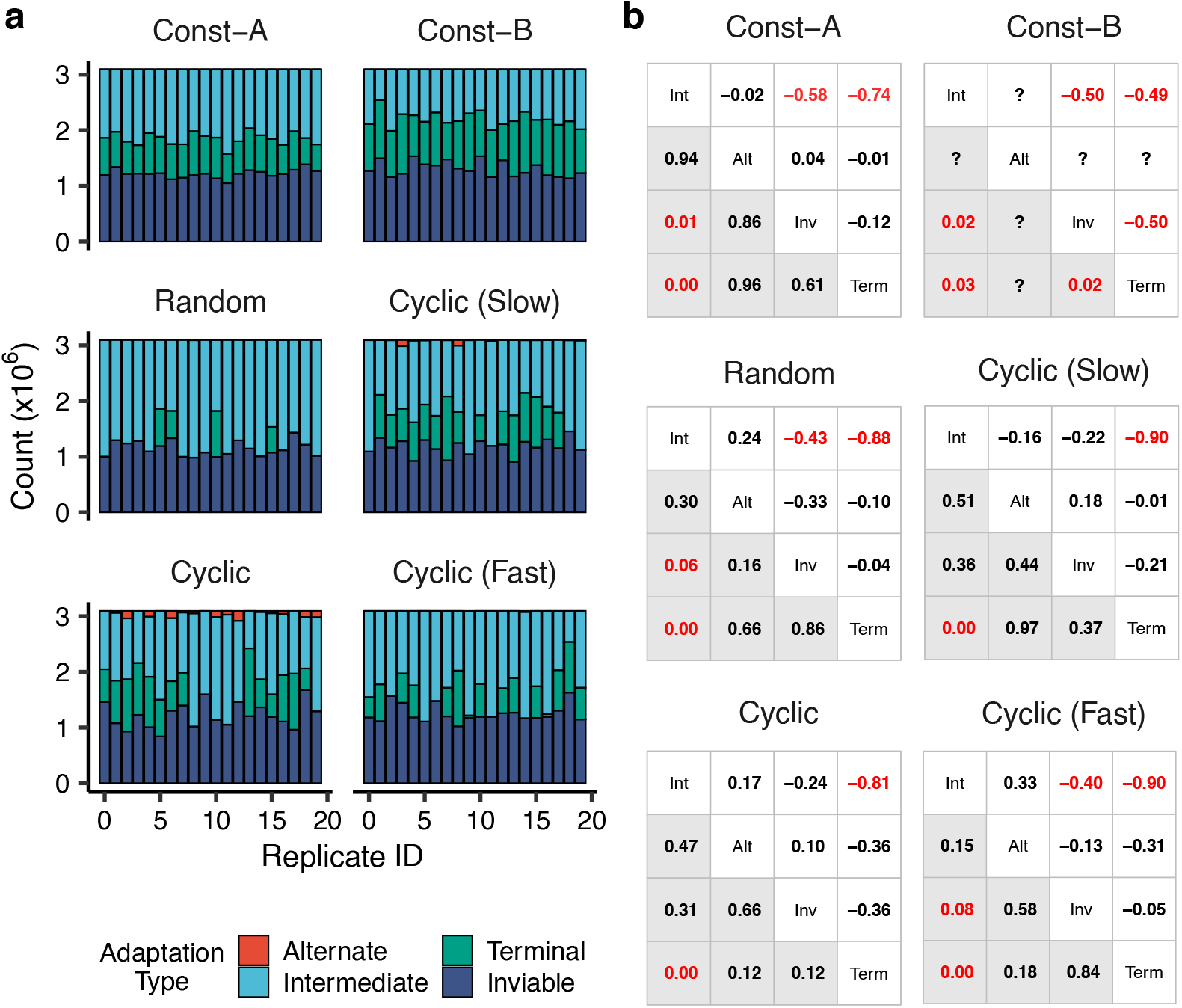
Distribution of different adaptations in the mutational neighborhood of dominant genotypes. (**a**) The number of mutants of the dominant genotype - isolated from populations evolved in the different regimes - that are either perfectly adapted to the terminal environment (’Terminal’), the environment alternate to the terminal (’Alternate’), have an intermediate phenotype (’Intermediate’) or are unable to self-replicate (’Inviable’). Each bar corresponds to one dominant genotype from a replicate population evolved under a particular environment change regime. Adaptations for both single and double mutants have been plotted here. (**b**) Correlation matrix between the number of mutants with different adaptation types for dominant genotypes from the six regimes. The upper diagonal numbers denote the Pearson correlation coefficient between the number of mutants with given adaptation types. The lower diagonal numbers (grey) denote the respective p-values for these correlations. Significant p-values (*<* 0.05) and respective correlation coefficients have been colored red. Text on the diagonal indicates the correlation being measured (Int=Intermediate, Alt=Alternate, Inv=Inviable, Term=Terminal).

**Figure S3:**
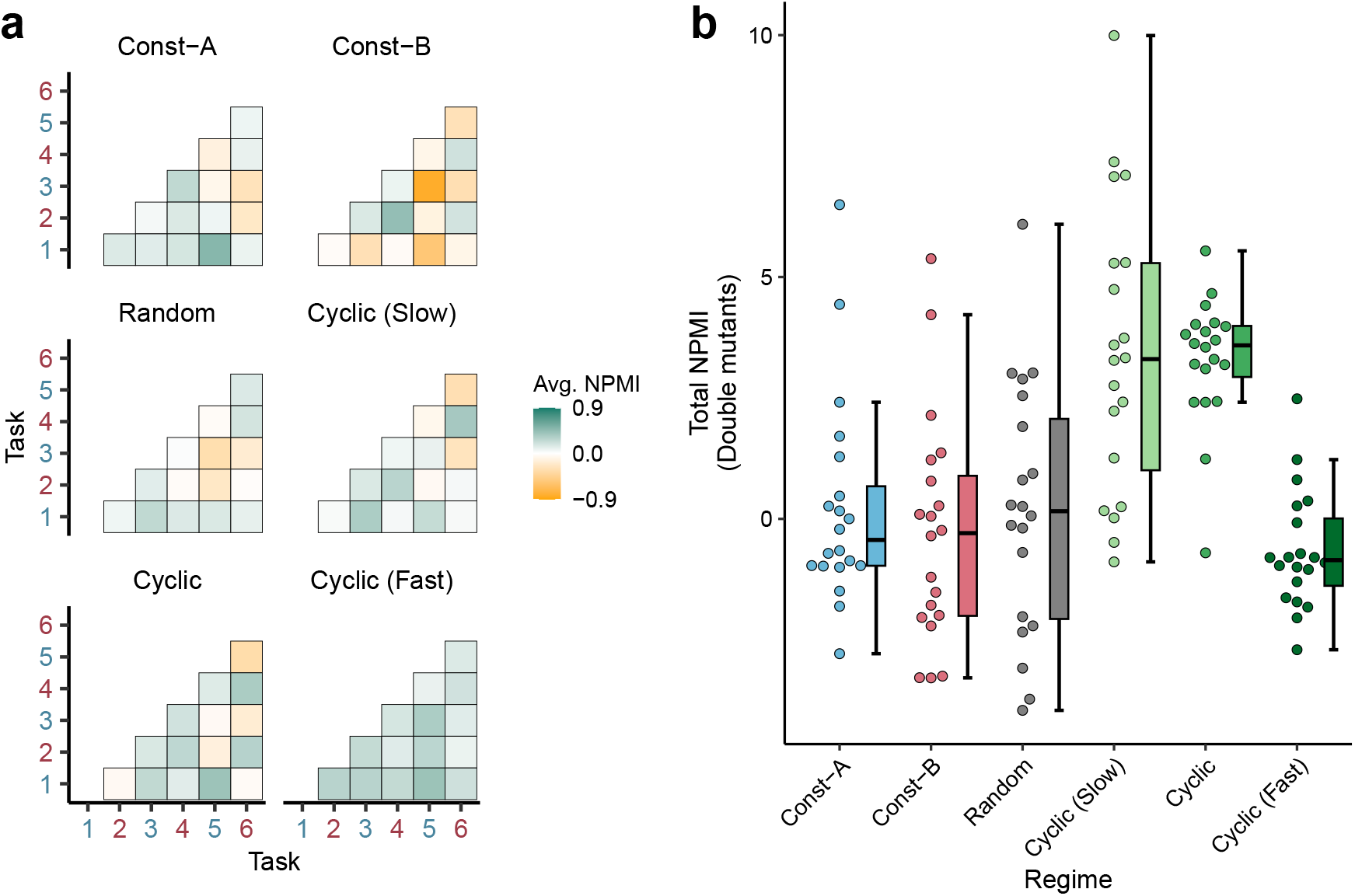
NPMIs evolved under fixed mutation rate experiments after removal of perfectly adapted mutants. (**a**) Pairwise NPMI between different task pairs plotted using double mutant data. Mutants that are perfectly adapted to environment A or B have been removed. Axes labels in blue and red denote tasks rewarded in environments A and B, respectively. (**b**) Total NPMIs calculated using the pairwise NPMIs in the previous panel. Total NPMIs are calculated by summing up the individual task-pair NPMIs after assigning them a positive or negative sign based on their cognate or non-cognate nature. Box-plot characteristics are the same as in Fig. 1.

**Figure S4:**
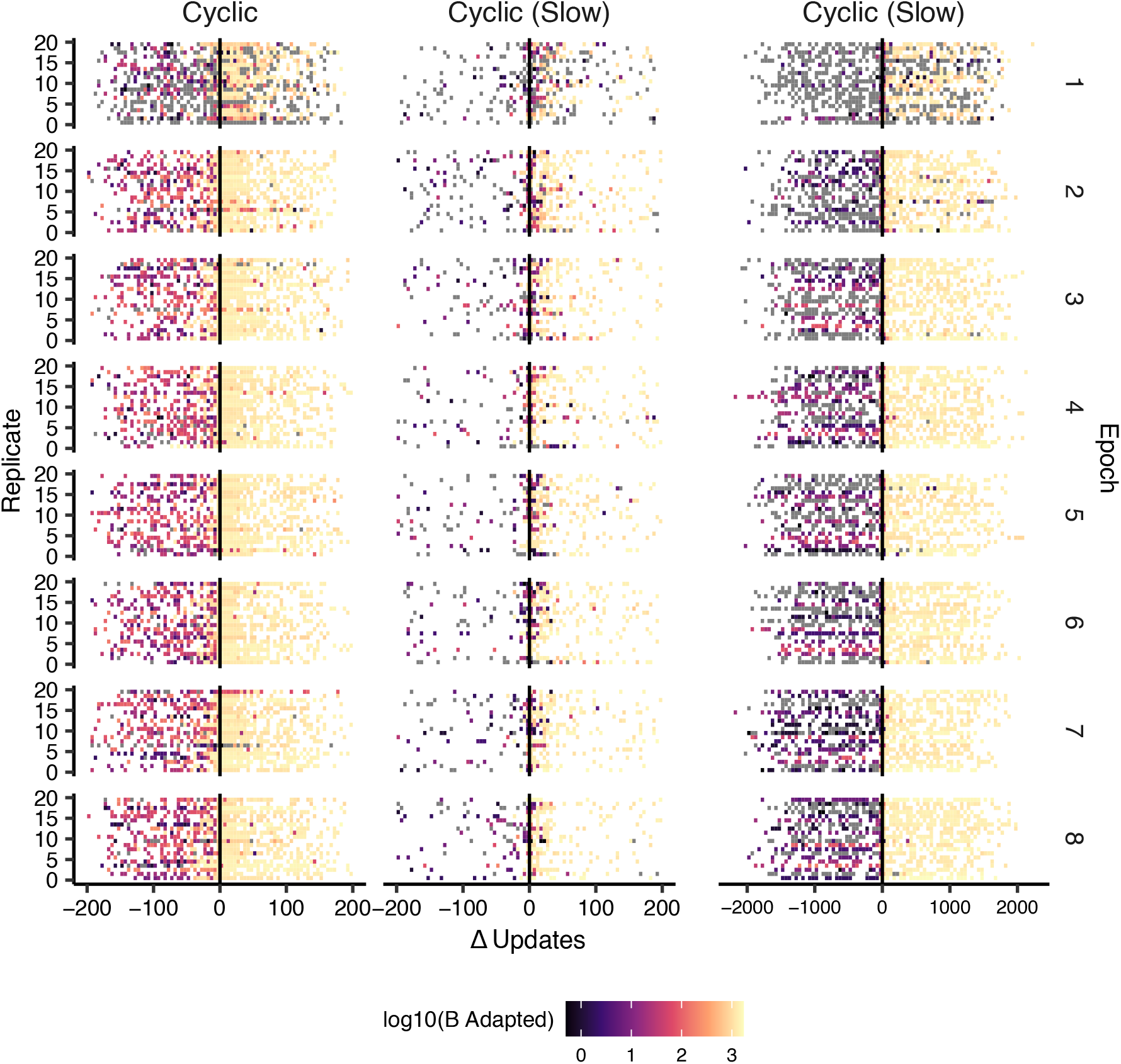
Median number of point-mutants of lineage sequences adapted to environment B before and after the environment switches from state A to B. The x-axis tracks the number of updates from the point of environmental transition where lineage mutants are recorded. The experiment time has been divided into eight epochs of 37,500 updates (approx. 375 generations) each, shown here as different rows. The y-axis in each figure shows 20 different replicates. Median tallies of point mutants were computed within intervals: bins of 5 updates for the two leftmost graphs, and an expanded bin size of 50 updates for the graph on the far right. The middle *Cyclic (Slow)* figure restricts the range to *±*200 updates.

**Figure S5:**
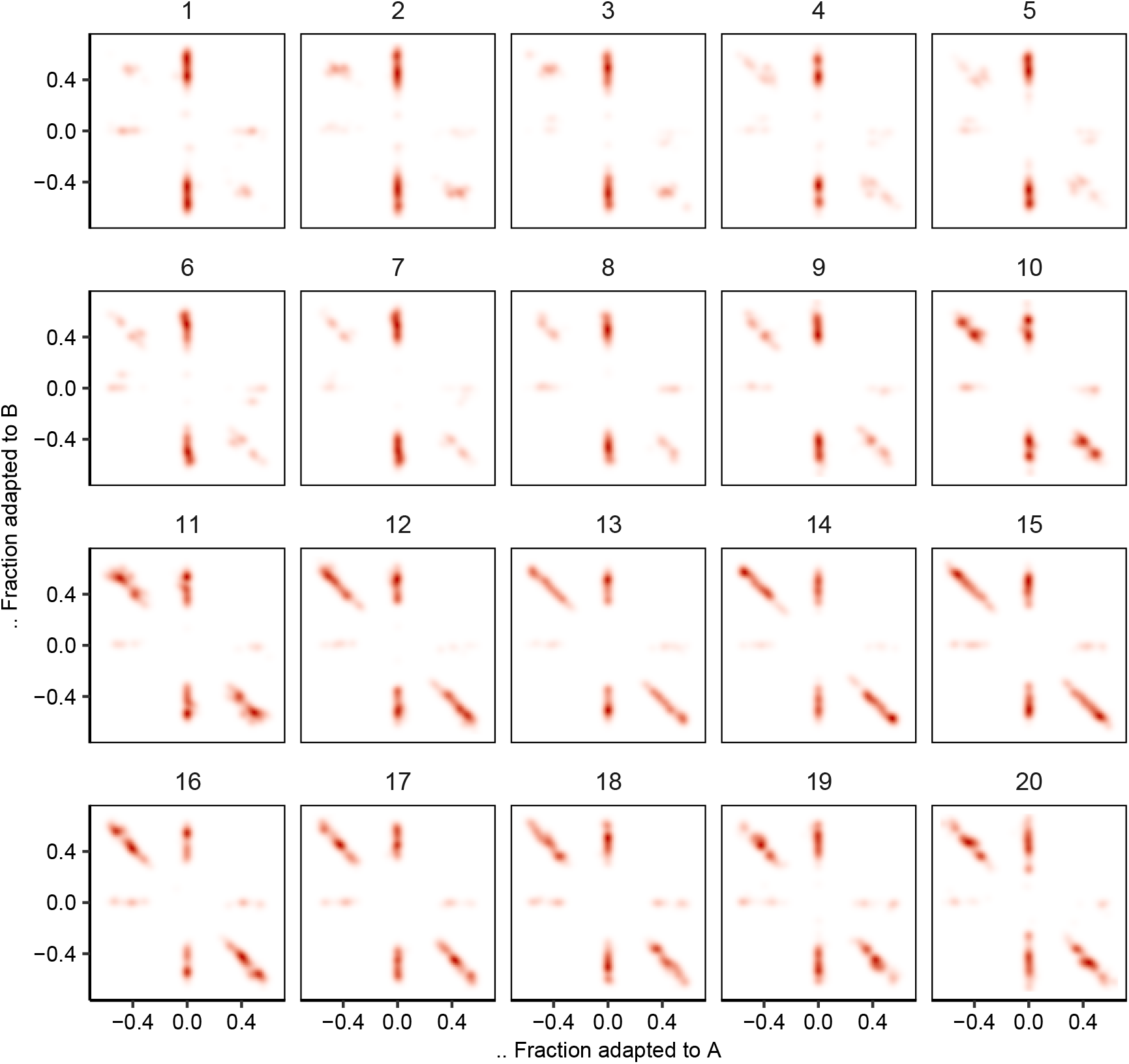
Mutational trade-offs over evolutionary time under the Cyclic regime. Density of points in the *trade-off plane* (See figure S1d and methods) plotted over 20 equally spaced time intervals during evolution in the *Cyclic* regime (all replicates). Each panel is a time interval spanning 15000 updates (approx. 1500 generations). Points along the *y* = *−x* diagonal represent mutation events on a lineage where an increase in number of mutants adapted to a particular environmental state was accompanied by an equal decrease in the number of mutants to the alternate environmental state.

**Figure S6:**
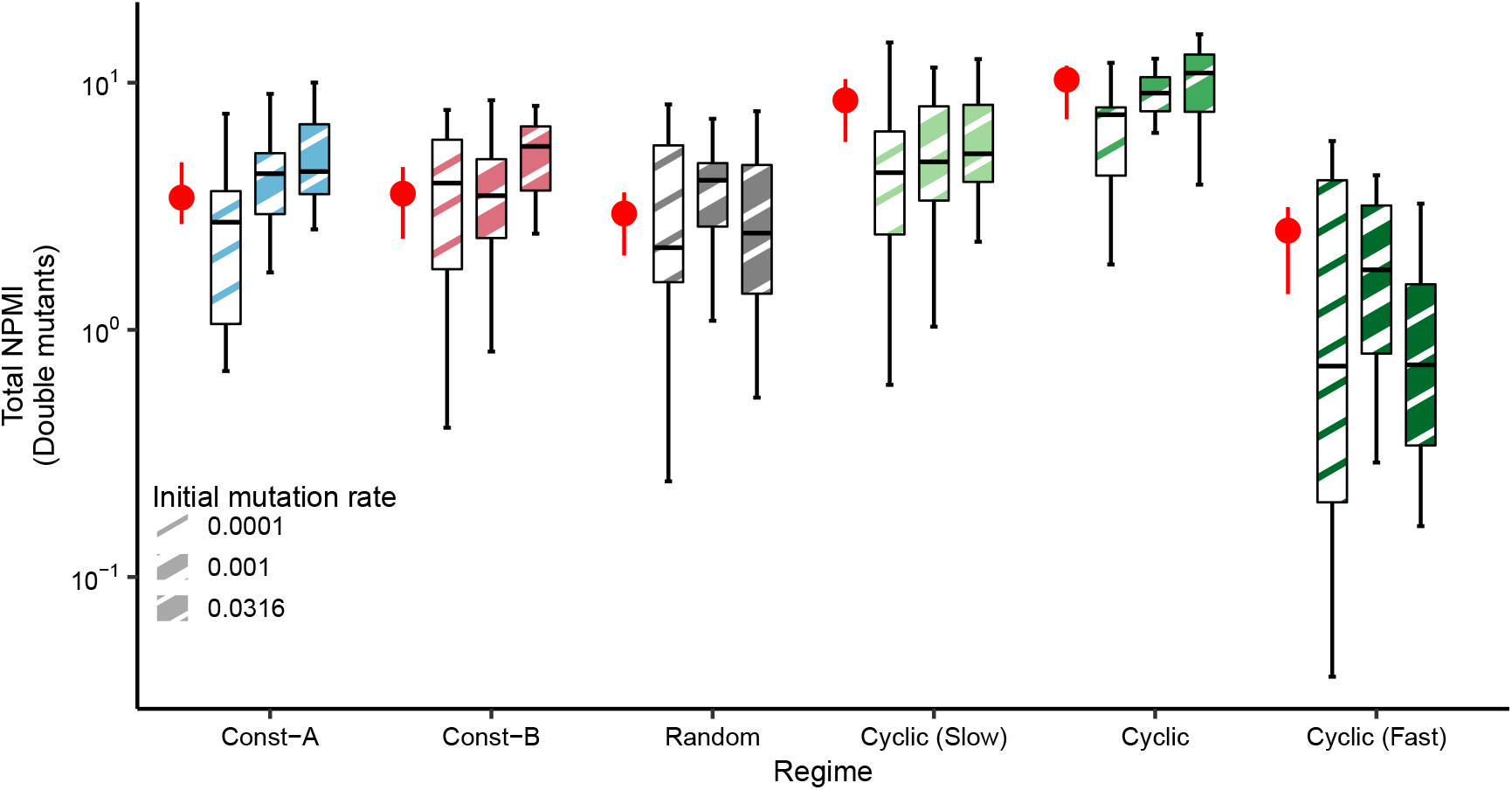
Evolved total NPMIs under fixed and evolving mutation rates. Total NPMI for the dominant genotypes isolated from the six environmental change regimes with evolving mutation rates. Box plots with different patterns denote NPMI values from experiments that started with different initial mutation rates. The red filled circles denote the median NPMI for genotypes from the fixed mutation rate experiments, with the error bars denoting the lower and upper quartiles.

**Figure S7:**
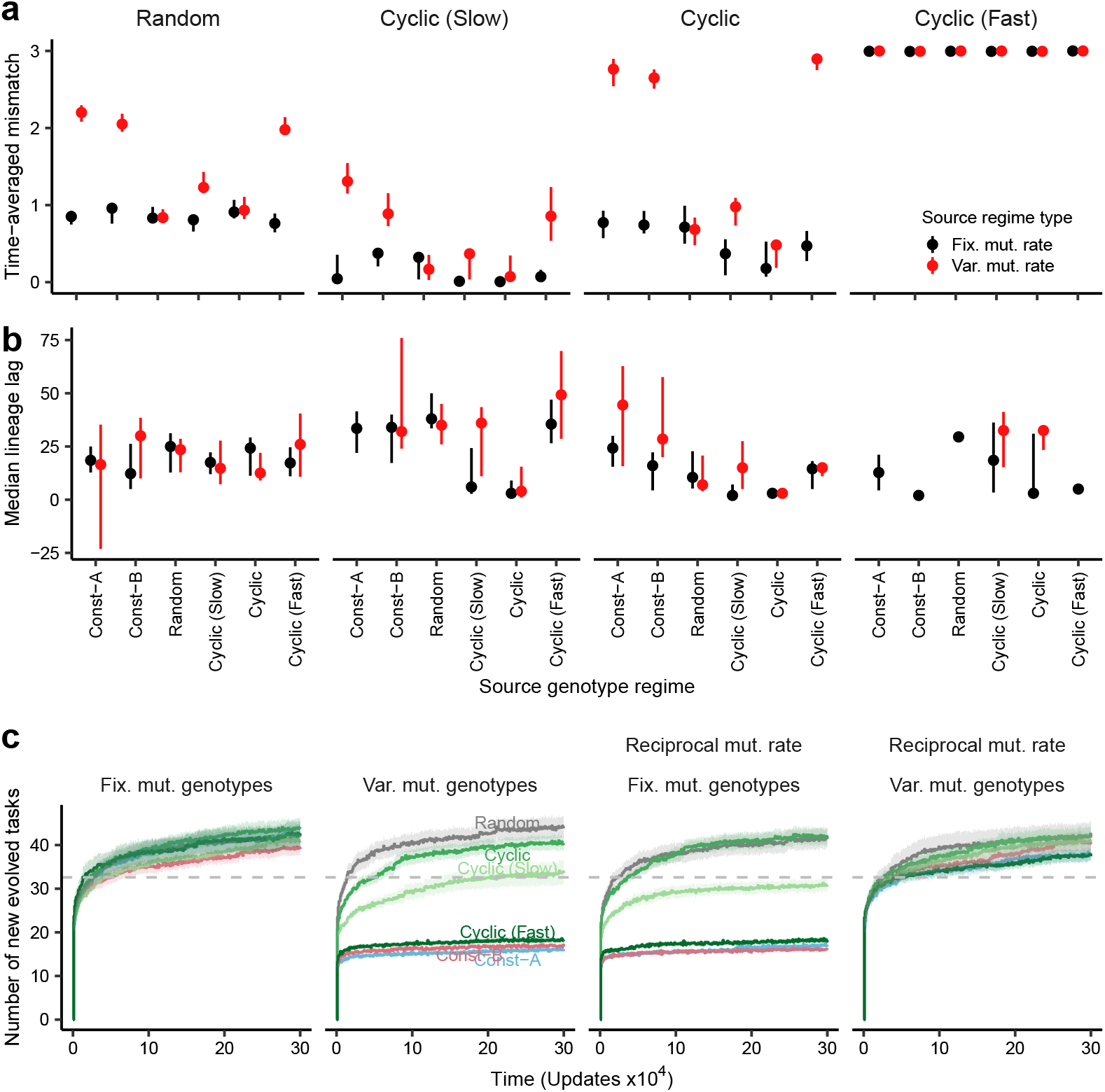
Performance of evolved genotypes transplanted into other regimes and completely new environments. (**a**) Median value of the time-averaged mismatch (over 10,000 updates) for genotypes isolated from primary evolution experiments in different test regimes (columns). The mismatch is calculated as the hamming distance between the dominant lineage phenotype and the environmental state—i.e, the number of task gains/losses required by the phenotype to be perfectly adapted to the extant environment. The x-axis denotes the regime the genotype was isolated from. (**b**) The median value of the mean lineage lag for genotypes from primary evolution in different test regimes. The lineage lag measures the time taken by the lineages to achieve a phenotype perfectly adapted to the environment after a switch. (**c**) The number of tasks evolved by genotypes from different source regimes when evolving in a new environment with 127 new tasks that increase fitness incrementally. The first panel shows the evolution of tasks in this new environment for genotypes isolated from the fixed mutation rate experiments (with a mutation rate of 10^−3^). The second panel shows the evolution of genotypes from evolving mutation rate experiments, retaining their evolved mutation rates. The third panel shows the evolution of fixed mutation rate genotypes when assigned the average evolved mutation rate from the evolving mutation rate experiments. The fourth panel shows the evolution of genotypes from evolving mutation rate experiments when assigned a basal mutation rate of 10^−3^. The dark lines denote the average across 20 replicates, and the ribbons represent the standard error.

**Figure S8:**
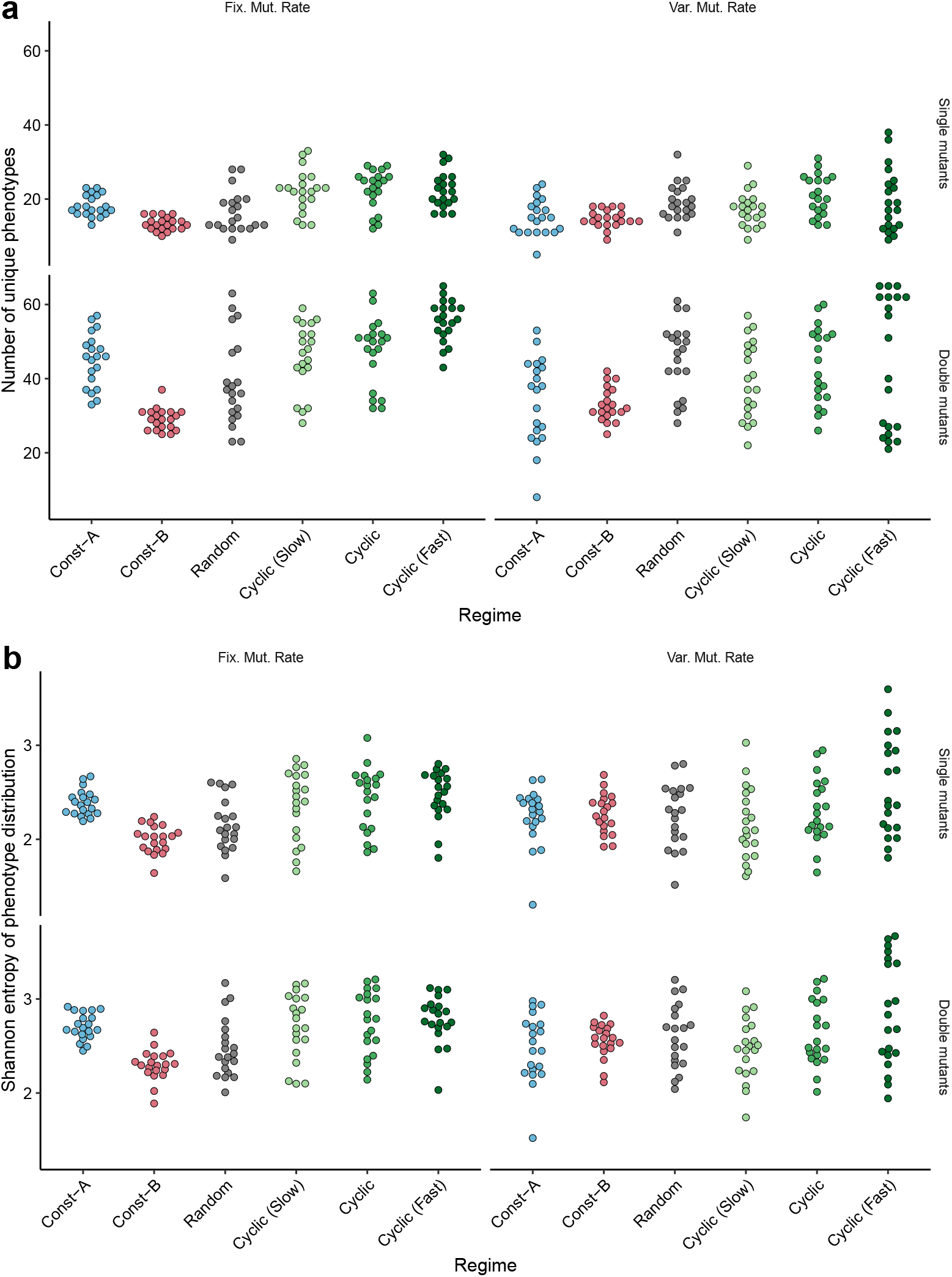
Diversity of phenotypes in the mutational neighborhood of evolved genotypes. (**a**) Total number of unique phenotypes found in the mutational neighborhood of dominant genotypes isolated from different evolutionary regimes. Individual columns are different mutation rate experiments (fixed or variable). Rows denote the mutant identity being tested (single mutants or double mutants). (**b**) Shannon entropy of the distribution of phenotypes in the mutational neighborhood of dominant genotypes isolated from different evolutionary regimes.

**Figure S9:**
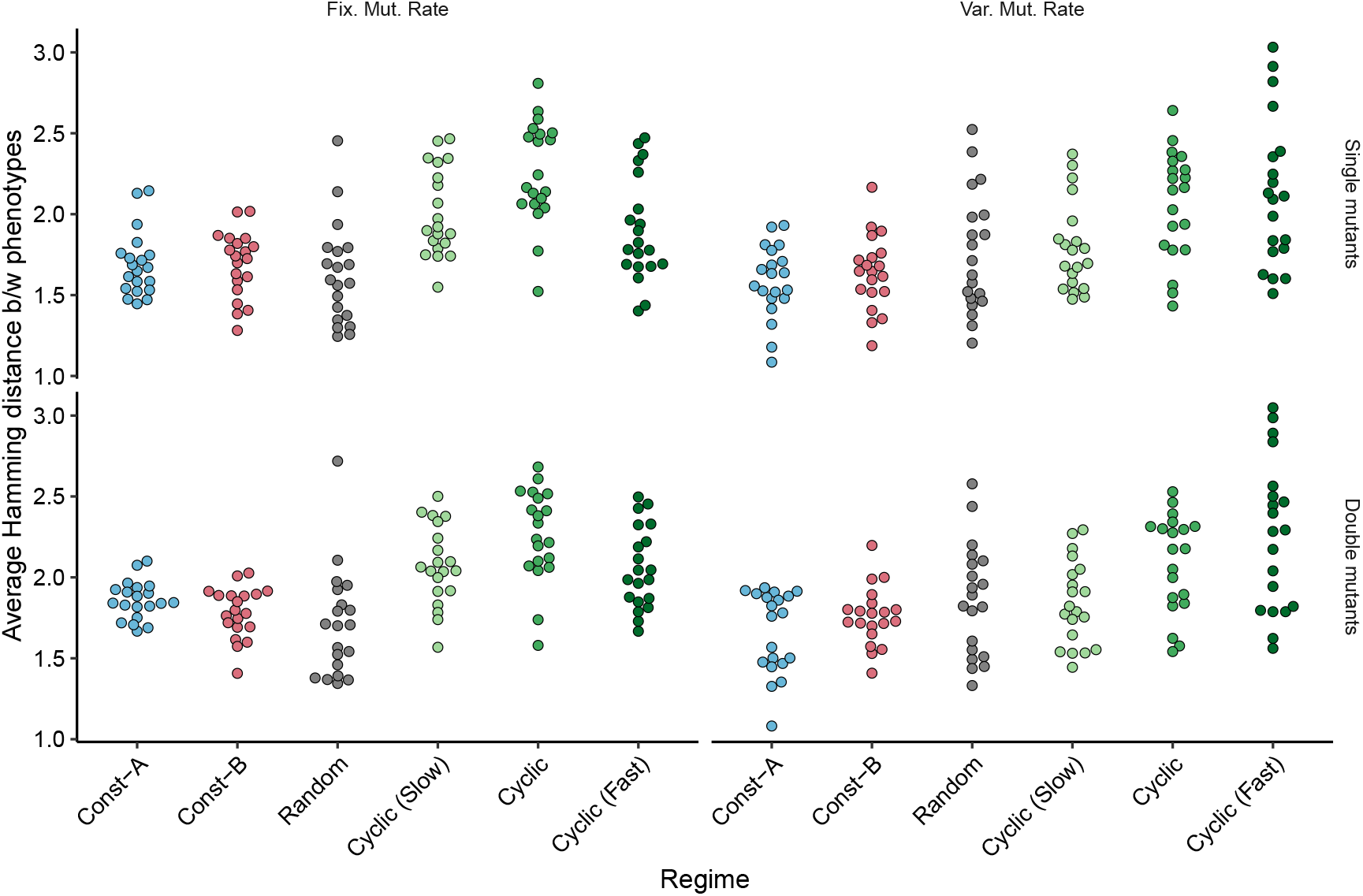
Phenotypic dissimilarity in the mutational neighborhood of evolved genotypes. Average hamming distance between phenotype pairs found in the mutational neighborhood of dominant genotypes isolated from different evolutionary regimes. Hamming distance is calculated as the number of task gains/losses required to convert one phenotype to another. Individual columns are different mutation rate experiments (fixed or variable). Rows denote the mutant identity being tested (single mutants or double mutants).

**Table 1:**
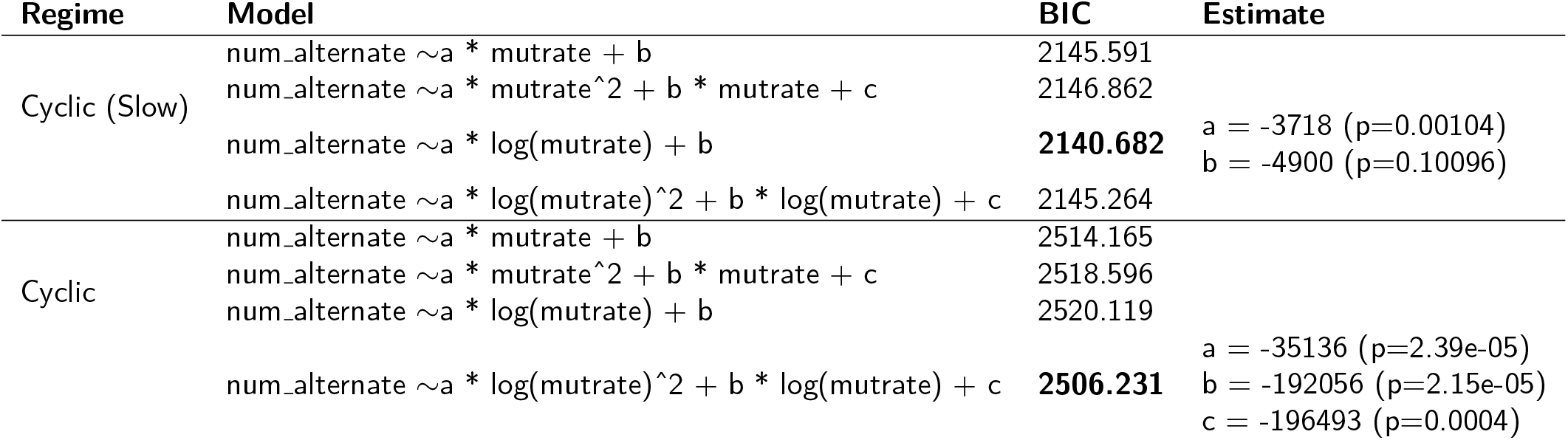
Statistical analyses associated with the number of alternate mutants evolved under different static mutation rates.

